# Targeting the cell membrane in established and emerging model organisms

**DOI:** 10.1101/2024.11.12.623055

**Authors:** Irene Karapidaki, Mette Handberg-Thorsager, Tsuyoshi Momose, Hitoyoshi Yasuo, Grigory Genikhovich, Sarah Assaf, Clara Deleau, Ying Pang, Clayton Pavlich, Beke Lohmann, Maria Lorenza Rusciano, Mattia Stranges, Juliette Mathieu, Marie Zilliox, Kirill Ustyantsev, Bastien Salmon, Béryl Laplace-Builhé, Manon Koenig, Jeffrey J. Colgren, Maria Ina Arnone, Eugene Berezikov, Thibaut Brunet, Gregor Bucher, Pawel Burkhardt, Daniel J. Dickinson, Evelyn Houliston, Jan Huisken, Lucas Leclère, Michalis Averof

## Abstract

Transgenic markers and tools have revolutionised how we study cells and developing organisms. Some of the elements needed to construct those tools are universally applicable (e.g. fluorescent proteins), while others are species-specific (e.g. cis-regulatory elements driving transcription). Membrane-localising signals that target proteins to the plasma membrane have been identified in several model organisms. Unfortunately, the efficacy of these signals varies greatly across species. To address this problem, we generated a toolkit of 11 membrane-localising tags that can be screened rapidly in diverse organisms. The toolkit includes tags that target the plasma membrane through different mechanisms, including signal peptides, the attachment of lipids, and fusion with lipid-binding domains. Each tag has been fused to the red fluorescent protein mScarlet3 and placed downstream of a T7 promoter, allowing the in vitro production of mRNA that can be readily delivered in a wide range of embryos and cells of interest. Through a collaborative effort, we tested this toolkit in ten species of animals spanning diverse phyla, including chordates, echinoderms, arthropods, nematodes, annelids, flatworms and cnidarians. We identify robust membrane-localising tags in each of these animals, and in one of animals’ closest relatives, the choanoflagellates. Three tags (KRas, GAP43 and Src64B) work in all of the species tested.

## Introduction

A range of localisation signals have been identified that direct proteins to specific parts of the cell. These are often peptide sequences that interact with the cells’ transport machinery, or direct chemical modifications that enable that targeting (von Heijne 1990; Casey 1995). The targeting peptides can often be fused with heterologous proteins, targeting them to specific cellular locations such as the nucleus, the plasma membrane or mitochondria. This approach has been used to localise a wide variety of marker and effector proteins in different parts of the cell, for live imaging, biochemical labelling, functional studies, gene editing, etc. Many targeting peptides function across eukaryotic species, which has helped to spread cellular markers and tools from established model organisms to new species of interest.

Various signals that target proteins to the plasma membrane have been identified and successfully applied across species (e.g. Greco et al. 2001; Onken et al. 2006; Prodon et al. 2010; Hadjieconomou et al. 2011; Kanca et al. 2013; Özpolat et al. 2017), but there is little consensus on which membrane-localising signals work best. As researchers studying non-conventional model organisms, we have found that several commonly used membrane-localising tags do not work sufficiently well in our species of interest.

A common problem is that membrane-tagged proteins do not localise exclusively to the plasma membrane, but also appear in endomembrane compartments, such as the endoplasmic reticulum, the Golgi, or endosomes. This is not surprising, as there is extensive trafficking between the plasma membrane and these compartments. The mechanisms that transport and anchor proteins selectively to the plasma membrane depend on interactions with other proteins and lipids in the plasma membrane. These may be influenced by signalling events, membrane composition and cell-cell contacts (e.g. Halet 2005; Gulyás et al. 2017; Morinaga et al. 2017), which can vary across species, developmental stages and cell types. Thus, there may not be a universal protein tag for targeting the plasma membrane in every species and cell type.

In this context, achieving plasma membrane localisation in new experimental systems requires testing of different tags to identify ones that work. Here, we present a toolkit of 11 membrane-localising tags, which can be readily tested in new species. These tags have been shown to drive heterologous proteins to the plasma membrane in at least some organisms. The sequences encoding these tags were fused with sequences encoding a red fluorescent protein, mScarlet3 (https://www.fpbase.org/protein/mscarlet3/; Gadella et al. 2023), and placed downstream of a T7 promoter to facilitate the *in vitro* production of mRNA. Once transcribed, these reporter mRNAs can be readily injected in embryos or cell types of interest.

The tags we include in this toolkit employ several different mechanisms to reach the plasma membrane. They include signal peptides for trafficking to the plasma membrane through the exocytic pathway, tags that direct the covalent attachment of different lipids, a domain that targets proteins to the plasma membrane independently of exocytosis or lipid modification, and lipid-binding domains that bind lipids which are predominantly located in the plasma membrane. The tags are summarised in Table 1 and described in more detail below.

**Table 1.** Plasma membrane-localising tags included in the toolkit.

| Name | Type of tag, mechanism | Previous use |
| --- | --- | --- |
| **HRas** | C-terminal prenylation (CAAX) and palmitoylation sites; attached on the cytoplasmic side of the plasma membrane | Mammalian cells (Hancock et al. 1991), *Drosophila* (Kanca et al. 2013), *Parhyale* neurons (Ramos et al. 2019), *Platynereis* (Özpolat et al. 2017), fission yeast (Onken et al. 2006) |
| **KRas** | C-terminal prenylation site (CAAX) and polybasic domain; attached on the cytoplasmic side of the plasma membrane | Mammalian cells (Hancock et al. 1991; Zhou et al. 2017), fission yeast (Haupt and Minc 2017) |
| **KRas6R** | Same as KRas, with 6K polybasic domain replaced by 6R; potentially interacting with different lipids | Mammalian cells (Hancock et al. 1991; Zhou et al. 2017) |
| **RitC** | C-terminal non-CAAX, non-exocytic membrane targeting domain; attached on the cytoplasmic side of the plasma membrane | Mammalian cells (Lee et al. 1996), fission yeast (Onken et al. 2006) |
| **GAP43** | N-terminal peptide carrying two palmitoylation sites; attached on the cytoplasmic side of the plasma membrane | *Ciona* (Roure et al. 2007), *Tribolium* (Benton et al. 2013), *Platynereis* (Lauri et al. 2014) |
| **Lyn** | N-terminal myristoylation and palmitoylation sites; attached on the cytoplasmic side of the plasma membrane | Mammalian cells (Gulyás et al. 2017), zebrafish (Haas and Gilmour 2006), *Drosophila* (Hadjieconomou et al. 2011), *Parhyale* (Alwes et al. 2016), *Platynereis* (Vopalensky et al. 2019) |
| **Src64B** | N-terminal myristoylation site from *Drosophila* Src64B; attached on the cytoplasmic side of the plasma membrane | *Drosophila* neurons (Pfeiffer et al. 2010) |
| **PH** | N-terminal mammalian PH domain, binding PtdIns(4,5)P2 lipids, enriched in the plasma membrane | Mammalian cells (Halet 2005), tunicates (Prodon et al. 2010; Guignard et al. 2020), *C.elegans* (Audhya et al. 2005; Heppert et al. 2016), hydrozoan embryos (Uveira et al. 2024) |
| **LactC2** | C-terminal phosphatidylserine (PS) binding domain, a lipid found in the inner leaflet of the plasma membrane | Mammalian cells (Yeung et al. 2008; Kay et al. 2012), budding yeast (Fairn et al. 2011), fission yeast (Haupt and Minc 2017) |
| **SP-CD8tm** | N-terminal signal peptide and transmembrane domain; trafficked in the exocytic pathway, attached on cytoplasmic side | *Drosophila* (Stapornwongkul et al. 2020) |
| **SP-GPI** | N-terminal signal peptide and C-terminal GPI attachment site; trafficked in the exocytic pathway, attached on extracellular side | Mammalian cells (Keller et al. 2001; Shelby et al. 2023), *Drosophila* (Greco et al. 2001) |

We evaluated the activity of these tags in a wide range of species, spanning seven animal phyla. These include both established and emerging model organisms belonging to the chordates (*Phallusia*), echinoderms (*Paracentrotus*), arthropods (*Parhyale*, *Tribolium*), nematodes (*Caenorhabditis*), annelids (*Platynereis*), flatworms (*Macrostomum*) and cnidarians (*Clytia*, *Pelagia*, *Nematostella*).

## Results and Discussion

### Selection of membrane-localising tags and reporter design

We built this toolkit by selecting tags that were previously shown to localise heterologous proteins to the plasma membrane, with preference for ones shown to work in different species. We were mindful to include tags that employ different mechanisms of localisation and/or different modes of attachment to the plasma membrane. The selected tags are summarised in Table 1 and Figure 1.

**Figure 1.**
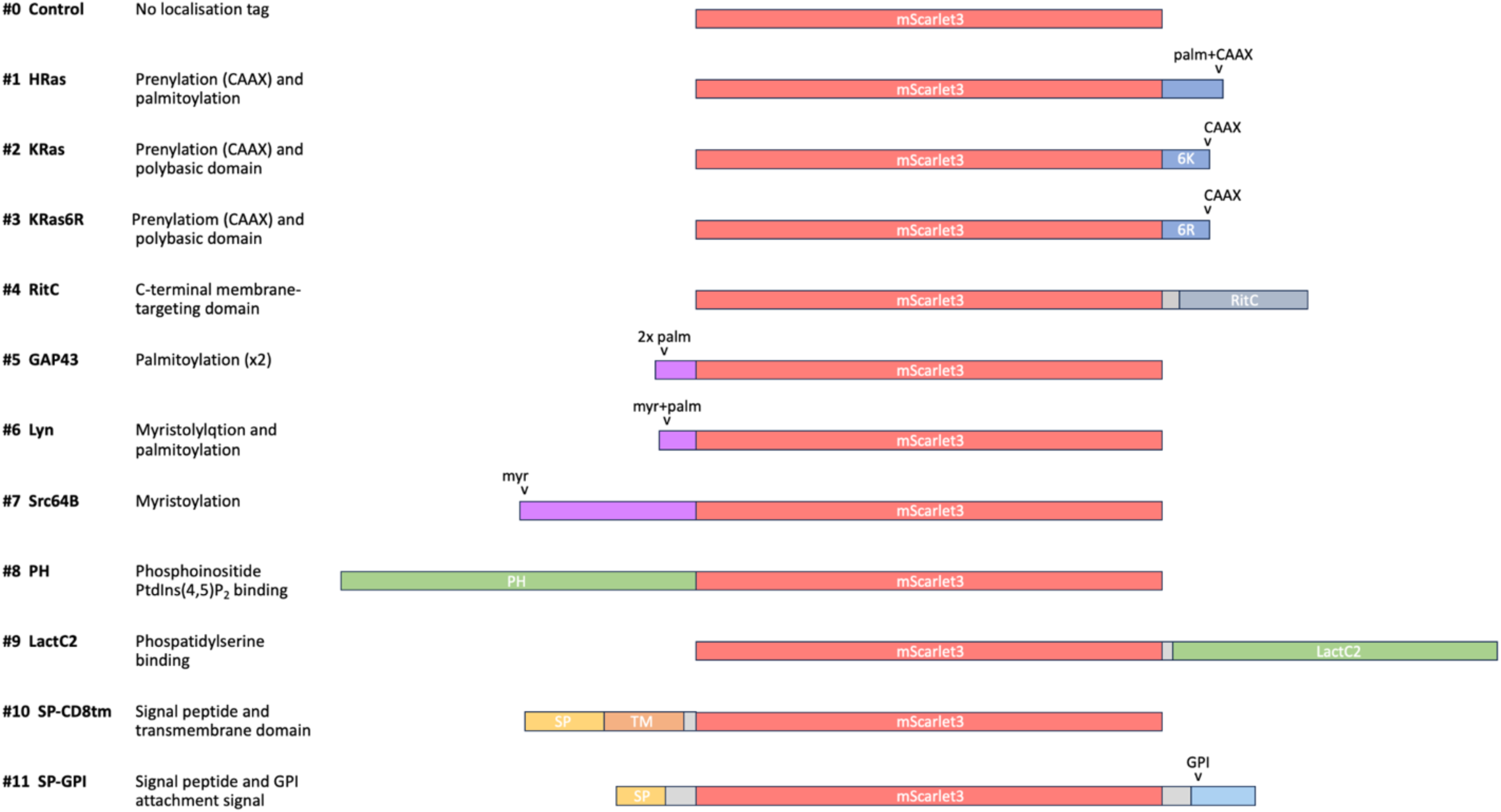
Toolkit of membrane-tagged constructs. Illustration of the eleven membrane-tagged constructs (#1-11) and the untagged control (#0), highlighting the mScarlet fluorescent protein shown (in red), C-terminal lipid addition tags (in blue), N-terminal lipid addition tags (in purple), lipid binding domains (in green), signal peptides (in yellow), transmembrane domain (in orange), a C-terminal GPI attachment tag (in cyan), and linkers (in grey).

One group of tags (constructs **#1 HRas**, **#2 KRas** and **#3 KRas6R**) includes C-terminal peptides that direct prenylation of the targeted proteins, i.e. the covalent attachment of isoprenoid lipids, typically geranyl or farnesyl lipid chains (Wang and Casey 2016). Prenylation tags typically carry a C-terminal CAAX motif and neighbouring sequences that may help to anchor the targeted protein to the plasma membrane. From this category, we include C-terminal peptides from the mammalian HRas and KRas proteins. The HRas tag carries a CAAX prenylation motif and a palmitoylation signal that together direct proteins to the plasma membrane through the exocytic pathway (Apolloni et al. 2000). The HRas tag has been used as a membrane-localising tag in mammalian cells and beyond (e.g. Kanca et al. 2013; Özpolat et al. 2017).

The tag from KRas isoform B (including exon 4B) carries a CAAX motif and a ‘polybasic’ domain, which includes positively-charged lysine residues that are thought to interact electrostatically with the negatively-charged lipids on the cytoplasmic side of the plasma membrane (Hancock et al. 1991; Apolloni et al. 2000; Gulyás et al. 2017). Previous studies have shown that a stretch of 6 lysine residues can be replaced by similarly charged arginine residues; the resulting variant KRas6R also drives robust localisation to the plasma membrane (Hancock et al. 1991). Interestingly, KRas and KRas6R appear to interact with different sets of lipids in the plasma membrane (Zhou et al. 2017), so we thought both should be included in this kit.

Besides the CAAX motifs of canonical Ras proteins, a C-terminal fragment of the mammalian Ras-like protein Rit (construct **#4 RitC**) has been shown to target proteins to the plasma membrane in the absence of a CAAX prenylation motif (Lee et al. 1996). Membrane targeting by RitC has been shown to work in mammalian cells and in fission yeast; it is thought to occur independently of the exocytic pathway and without a post-translational attachment of lipids (Onken et al. 2006).

Another group of tags (constructs **#5 GAP43**, **#6 Lyn** and **#7 Src64B**) includes N-terminal peptides that direct the myristoylation and/or palmitoylation of the targeted proteins, i.e. the covalent attachment myristic or palmitic fatty acids (Resh 1999), which direct targeting to the exocytic pathway and the plasma membrane. C-terminal myristoylation and palmitoylation tags have been used to label plasma membranes in vertebrates, tunicates, annelids and arthropods (Haas and Gilmour 2006; Roure et al. 2007; Pfeiffer et al. 2010; Hadjieconomou et al. 2011; Benton et al. 2013; Lauri et al. 2014; Alwes et al. 2016; Vopalensky et al. 2019).

Two tags (constructs **#8 PH** and **#9 LactC2**) consist of protein lipid-binding domains, which bind lipids that are enriched in the plasma membrane. The N-terminal PH domain of human PLCdelta1 has a strong affinity and specificity for PtdIns(4,5)P2, a phosphoinositide that resides primarily in the plasma membrane (Halet 2005). It has been used to label cell membranes in tunicate, *C. elegans* and hydrozoan embryos (Audhya et al. 2005; Prodon et al. 2010; Heppert et al. 2016; Guignard et al. 2020; Uveira et al. 2024). The C-terminal domain of bovine Lactadherin (LactC2) binds specifically phosphatidylserine (PS), a phospholipid that is enriched in the inner leaflet of plasma membranes (Yeung et al. 2008). LactC2 has been used as a sensor of PS in the plasma membranes of mammalian cells, budding and fission yeast (Yeung et al. 2008; Fairn et al. 2011; Kay et al. 2012; Haupt and Minc 2017).

Our toolkit also includes two constructs carrying signal peptides, expected to reach the plasma membrane through canonical exocytosis via the endoplasmic reticulum and the Golgi. One of these constructs carries the transmembrane domain of the mammalian CD8 protein, placing the tagged fluorescent on the cytoplasmic side of the plasma membrane (construct **#10 SP-CD8tm**); it has previously been used as a membrane-localising tag in *Drosophila* (Stapornwongkul et al. 2020; of note, this construct differs from *Drosophila* constructs that contain full-length CD8, see Lee and Luo 1999; Hadjieconomou et al. 2011; Rujano et al. 2022). The second construct carries a glycosylphosphatidylinositol (GPI) attachment signal (construct **#11 SP-GPI**); during transit through the endoplasmic reticulum, GPI is covalently attached at this site (Orlean and Menon 2007; Kinoshita 2020). Unlike the other membrane anchoring tags, the GPI anchor tethers the fluorescent protein on the extracellular side of the plasma membrane. The GPI tag we selected has been used previously in *Drosophila* and mammalian cells (Greco et al. 2001; Keller et al. 2001).

In addition to these tagged constructs, we generated a negative control in which the mScarlet3 is expressed without a localisation tag (construct **#0 Control**).

Each of these tags was placed at the N- or C-terminus (see Table 1) of the mScarlet3 fluorescent protein, flanked with 5’ and 3’ untranslated (UTR) sequences and placed downstream of the T7 promoter (see Materials and Methods). The plasmids carrying those constructs were used as templates for *in vitro* transcription, to produce the corresponding mRNAs.

### Screening the membrane tags in diverse animal species

We tested these membrane tags by microinjecting each mRNA in the eggs, embryos or gonads of ten species – *Phallusia mammillata*, *Paracentrotus lividus, Parhyale hawaiensis*, *Tribolium castaneum*, *Caenorhabditis elegans*, *Platynereis dumerilii*, *Macrostomum lignano*, *Clytia hemisphaerica*, *Pelagia noctiluca*, and *Nematostella vectensis* – using microinjection techniques specific to each organism (see Methods). The injected animals were then screened for reporter fluorescence (or immunofluorescence in the case of *Nematostella*) at specific stages of embryonic or post-embryonic development, and 3D image stacks were captured by confocal or light sheet microscopy. Results from each species are presented in Figures 2-5 and Suppl. Figures S1-S13.

**Figure 2.**
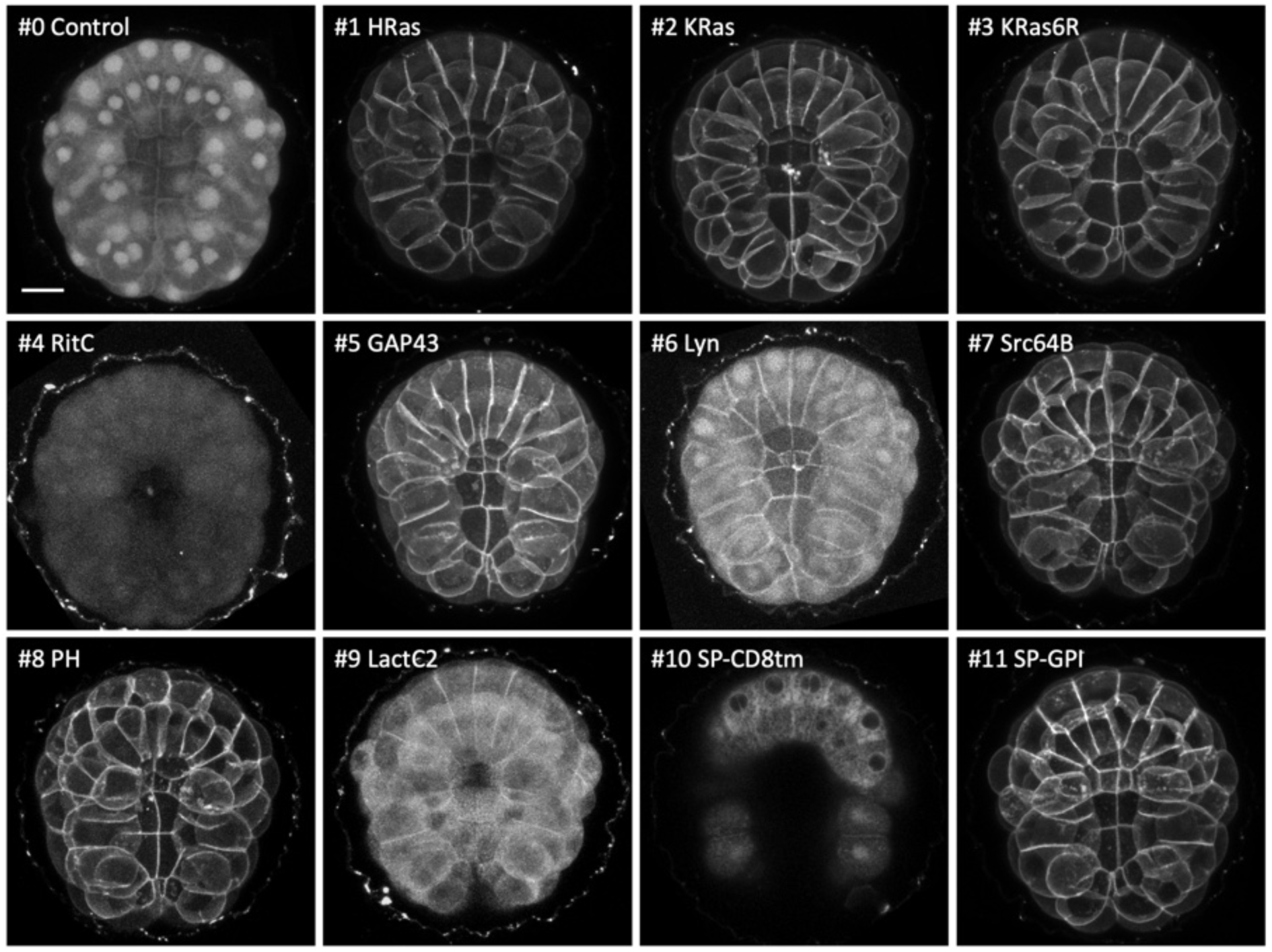
Localisation of membrane-tagged reporters in the tunicate *Phallusia mammillata*. mScarlet3 fluorescence in gastrulating *Phallusia* embryos, injected with mRNA of the membrane-tagged and control constructs, viewed from the vegetal pole. Images are maximum intensity projections, except for SP-CD8tm, which is a projection of three z-slices to highlight the perinuclear signal. Images for #4, #6, #8 and #9 were adjusted in brightness and contrast to reveal weak fluorescence. Scale bar, 25 μm.

**Figure 3.**
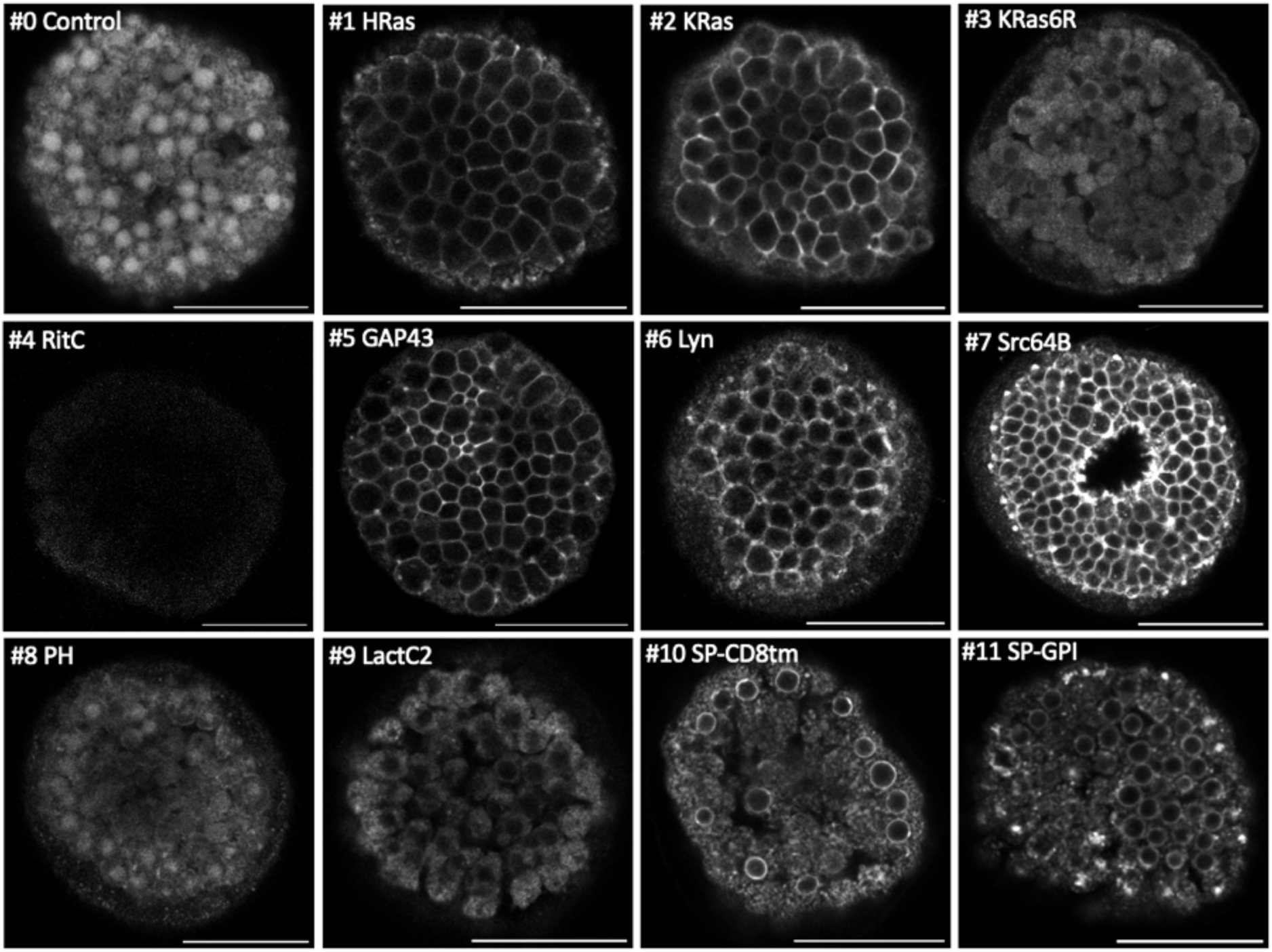
Localisation of membrane-tagged reporters in the sea urchin *Paracentrotus lividus*. mScarlet3 fluorescence in *Paracentrotus* blastula stage embryos (all except #7) or gastrula stage embryos (#7). Images are single confocal planes on the embryo’s surface. Some images were acquired with different settings (see Methods) and adjusted in brightness and contrast to reveal weak fluorescence. Scale bar, 50 µm.

**Figure 4.**
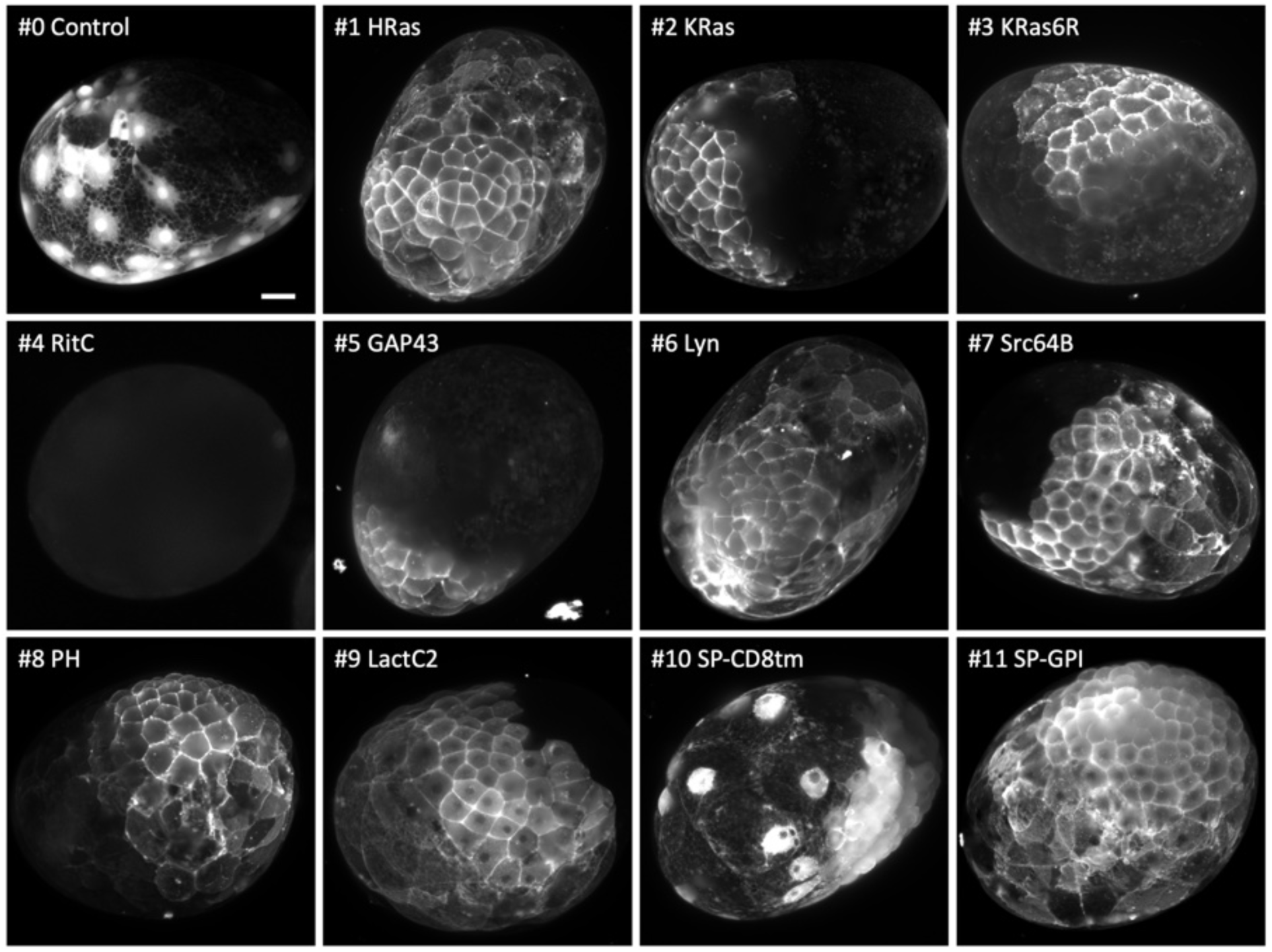
Localisation of membrane-tagged reporters in the crustacean *Parhyale hawaiensis*. mScarlet3 fluorescence in 1-day old *Parhyale* embryos, injected with mRNA of the membrane-tagged and control constructs. Only parts of each embryo express the reporter, due to uneven distribution of the injected mRNA. Images show maximum projections from light sheet microscopy captured with different exposure times (except #4, which gave no signal above background and was captured by conventional fluorescence microscopy). Scale bar, 50 µm.

**Figure 5.**
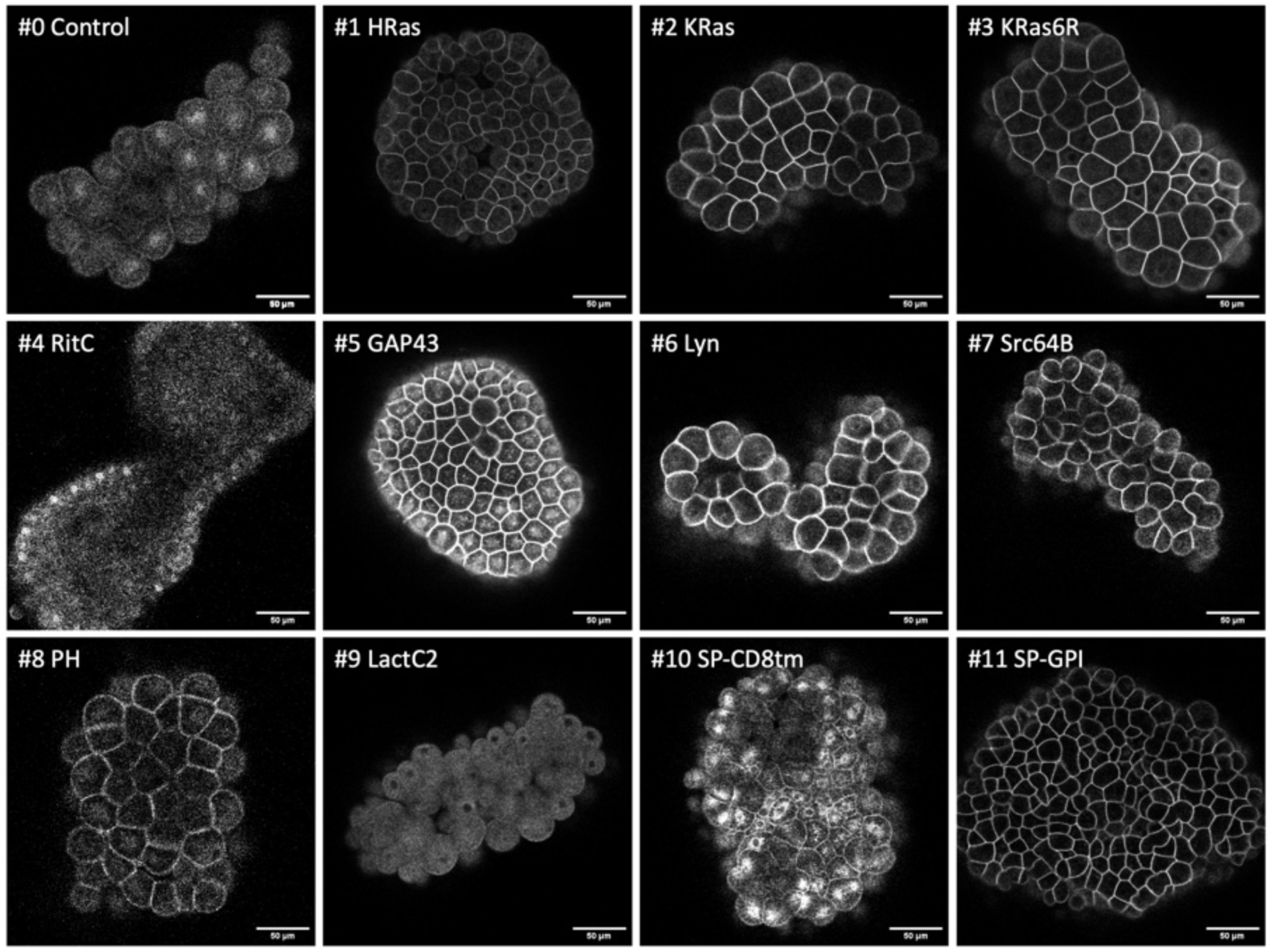
Localisation of membrane-tagged reporters in hydrozoan *Clytia hemisphaerica* embryos. mScarlet3 fluorescence in *Clytia* blastula-stage embryos, 5-6 hours after mRNA injection into oocytes. Images are single confocal planes of the epithelial surface. Images for #0, #4, #6, #7, #8 and #10 were adjusted in brightness and contrast to reveal weak fluorescence. Scale bar, 50 µm.

Multiple constructs showed significantly stronger fluorescence at the plasma membrane than at the interior of cells. While we could observe differences in fluorescence intensity among embryos injected with the same construct (presumably due to variation in the amount of mRNA injected and its spread within embryos), clear differences in the intensity and degree of localisation at the membrane could be observed between constructs (summarised in Figure 6). In *Phallusia, Paracentrotus, Parhyale*, *Macrostomum* and *Clytia* embryos, the fluorescence signals at the plasma membrane and in the interior of cells could also be quantified by image analysis (Suppl. Figures S1, S2, S4, S8 and S9).

**Figure 6.**
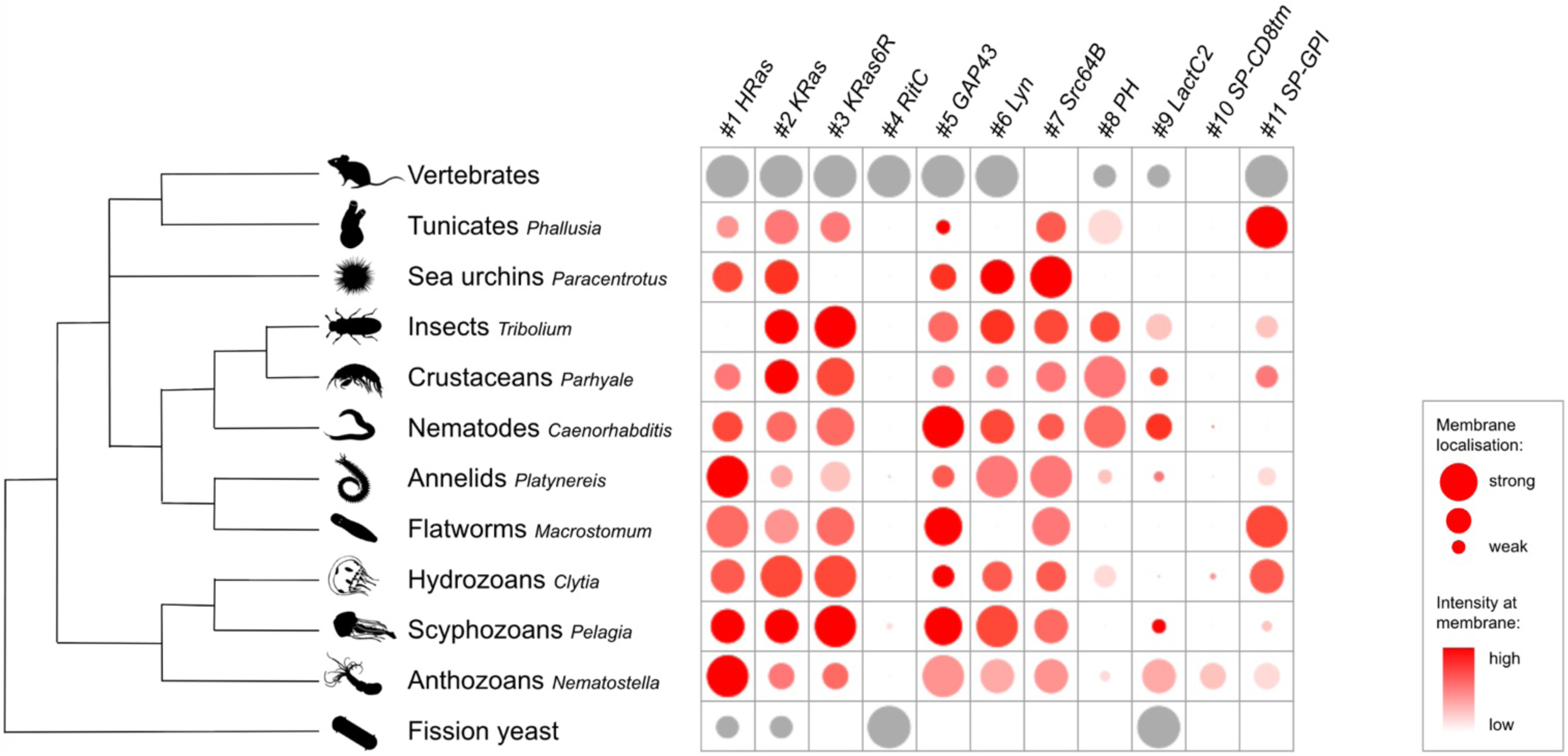
Overview of results from 11 membrane-localising tags across different organisms. For each species, relative values were assigned to each reporter indicating the degree of plasma membrane localisation (circle size) and fluorescence intensity at the membrane (colour intensity) (see Methods). The intensity estimates should be taken only as rough indications, as the quantity of mRNA delivered per embryo can vary considerably. Red circles represent data generated in this work; grey circles represent data from previous publications (see Table 1).

In most cases fluorescent protein localisation was scored within 24 hours from delivering the mRNA, but in some species, significant levels of fluorescence could also be observed a few days after mRNA injection. In *Platynereis*, fluorescence persisted in larval stages – particularly in the nervous system – and in some cases it was visibly localised to the plasma membrane (Suppl. Figure S7). In *Pelagia* and *Clytia*, membrane localisation could be scored in planula larvae and primary polyps, respectively, 2 and 3 days after fertilisation (Suppl. Figure S10).

The time required to screen the entire toolkit varies between species, depending on the availability of embryos, ease of injection, and method of screening. In some species (e.g. *Paracentrotus*, *Parhyale*, *Caenorhabditis*), once the conditions for performing the experiment are established, the entire set of constructs can be injected and screened by one person within a few working days.

### Comparing membrane localisation between tags and across species

Several tags in our toolkit were efficient in localising mScarlet3 to the plasma membrane in a wide range of animals (Figure 6). The C-terminal prenylation tags gave excellent membrane localisation in most species, although the efficacy of the three constructs (HRas, KRas and KRas6R) differed in some species. Some of the N-terminal lipidation tags, particularly Src64B, also gave excellent membrane localisation. GAP43 and Lyn localised to the plasma membrane in most species, but not as robustly as Src64B.

The phosphoinositide-binding PH domain showed specific membrane fluorescence in *Phallusia*, *Tribolium*, *Parhyale* and *Caenorhabditis*, and very weak fluorescence at the plasma membrane in *Clytia*. For many years, the PH domain has been used as the default membrane marker in *Phallusia* and *Caenorhabditis* (Prodon et al. 2010; Guignard et al. 2020, Audhya et al. 2005; Heppert et al. 2016), but we find here that it is more weakly expressed compared to other markers. In *Clytia* the weak signal can be considerably improved by using endogenous UTRs and codon optimised mCherry (Suppl. Figure S15).

The phosphatidylserine-binding LactC2 tag gave variable fluorescence intensity and membrane localisation in different species. In *Parhyale*, it gave strong fluorescence in the plasma membrane and weaker fluorescence in the cytoplasm, delineating cell boundaries, cytoplasm and nuclei very clearly.

The SP-CD8 tag, containing a signal peptide and the transmembrane domain of CD8, which has previously been used in *Drosophila* (Stapornwongkul et al. 2020), did not robustly localise to the plasma membrane in any of the species we tested. Live imaging in *Parhyale* embryos, shows SP-CD8tm-mScarlet3 localising transiently in the plasma membrane, but most of the fluorescence ends at the interior of the cell (Suppl. Video 1). The SP-GPI tag localised mScarlet3 very robustly to the plasma membrane in *Phallusia*, *Macrostomum* and *Clytia*, but its efficacy varied widely across species.

The C-terminal RitC domain, which has worked well in mammalian cells and in fission yeast (Lee et al. 1996; Onken et al. 2006), showed no or very little detectable fluorescence in all the species. This could be due to inefficient translation, misfolding or instability of the fusion protein.

By screening the toolkit presented here we identify three membrane-localising tags that were effective in most tested species: KRas, GAP43 and Src64B (Figure 6). Among these, GAP43 gave a somewhat lower membrane-to-cytoplasm signal. Another six tags, namely HRas, KRas6R, Lyn, PH, LactC2 and SP-GPI, performed well in some species but not in others. Two tags, RitC and SP-CD8tm, failed in most species.

From a species perspective, we identified several efficient membrane-localising tags in *Paracentrotus*, *Parhyale*, *Macrostomum*, *Clytia* and *Pelagia*, where such tags were not available until now, and expanded the range of tags available in *Phallusia*, *Tribolium*, *Platynereis, Caenorhabditis* and *Nematostella*.

Given the wide phylogenetic distribution of the animals that we tested, we expect these results will serve as a useful guide for selecting effective membrane-localising tags in a wide range of animals, and perhaps more widely in phylogeny. In this context, we tested one of the tags that shows conserved activity across the animals, KRas (#2), in the chanoflagellate *Salpingoeca rosetta*. Choanoflagellates are the closest living relatives of the animals, comprising unicellular and colonial forms (Brunet and King 2017). We transfected *S. rosetta* with a plasmid expressing the mStayGold fluorescent protein tagged with the KRas membrane-localising sequence. We found that the KRas-tagged fluorescent protein localises robustly to the plasma membrane, including the microvilli of the collar and the flagellum (Figure 7).

**Figure 7.**
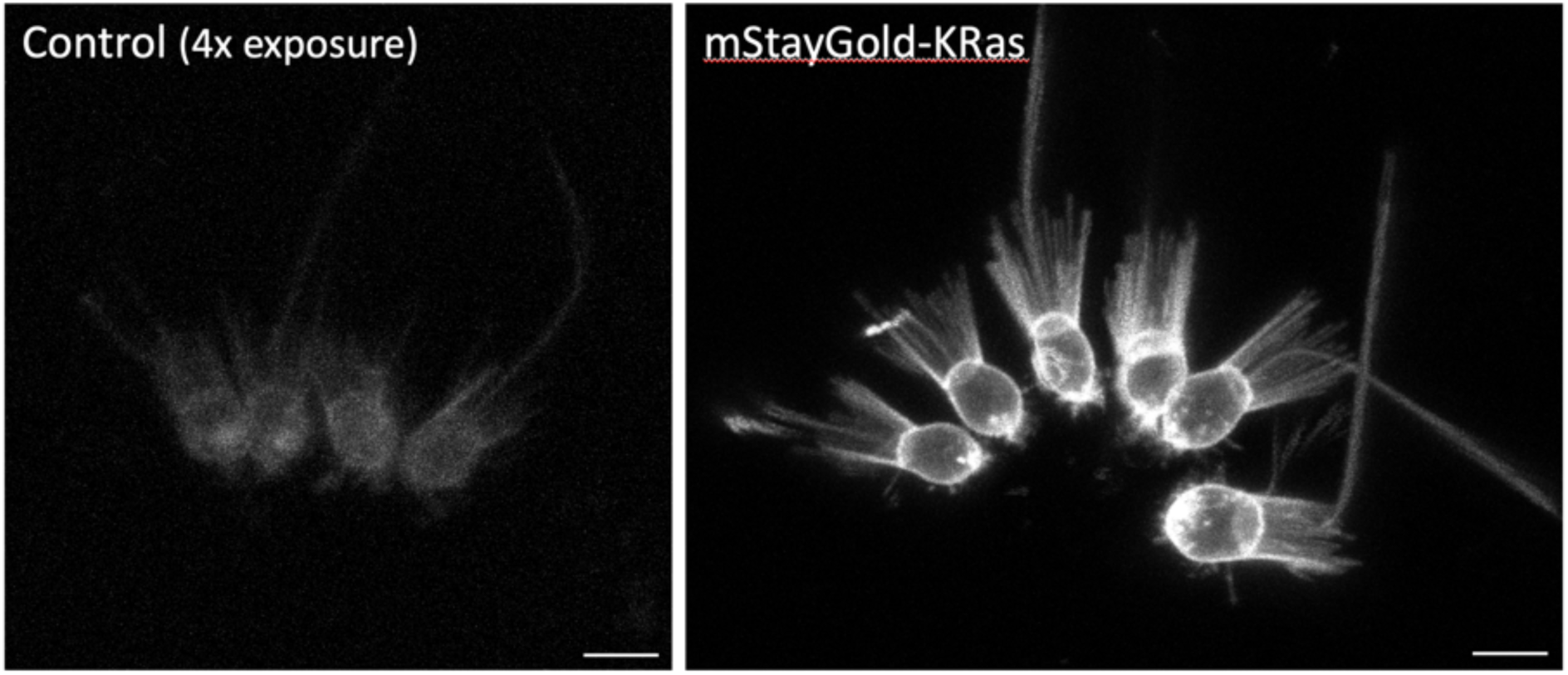
Localisation of a membrane-tagged reporter in the choanoflagellate *Saplingoeca rosetta*. *S. rosetta* cells transfected with an mStayGold-KRas expressing plasmid (right), compared with untransfected cells (left, shown with a 4-fold increase in brightness to reveal background levels of fluorescence). KRas-mStayGold localises to the plasma membrane, including the microvilli of the collar and the flagellum. The images show maximum projections of confocal stacks. Scale bars, 5 µm.

### Limitations and further testing

Protein localisation to the plasma membrane can vary not only between species, but also between cell types and cell states within a species. Such intra-species differences may be due to variations in the way protein trafficking is organised in different cell types (e.g. in polarised epithelial versus unpolarised mesenchymal cells), variations in membrane lipid composition, or differences in cell-cell contracts, signalling, and protein or lipid modifications that affect trafficking or anchoring to the plasma membrane (e.g. Halet 2005; Carvalho et al. 2012; Morinaga et al. 2017). Membrane localisation in one cell type does not necessarily predict localisation in another. Screening our toolkit by microinjecting early embryos therefore serves as a rapid first screen, from which promising candidates can be selected for further testing in other developmental stages and cell types of interest.

In these follow-up experiments, the selected membrane tags can be combined with different fluorescent proteins and incorporate 5’ and 3’ UTRs from the target species to improve expression. To provide an example, in *Clytia* we found that fluorescence could be enhanced several fold by using endogenous UTRs and codon optimised mCherry (Suppl. Figure S15).

In *Parhyale*, we have used two of the membrane-localising tags that we identified here to generate transgenic reporter lines. Specifically, the KRas and Src64B tags were combined with the mNeonGreen fluorescent protein, cloned downstream of a heat-inducible promoter and stably inserted in the *Parhyale* genome (see Methods). These transgenic reporters allow us to visualise the shape and behaviour of epithelial, muscle and neuronal cells in living embryos, juveniles and adults, including cell dynamics during the course of leg regeneration (Figure 8).

**Figure 8.**
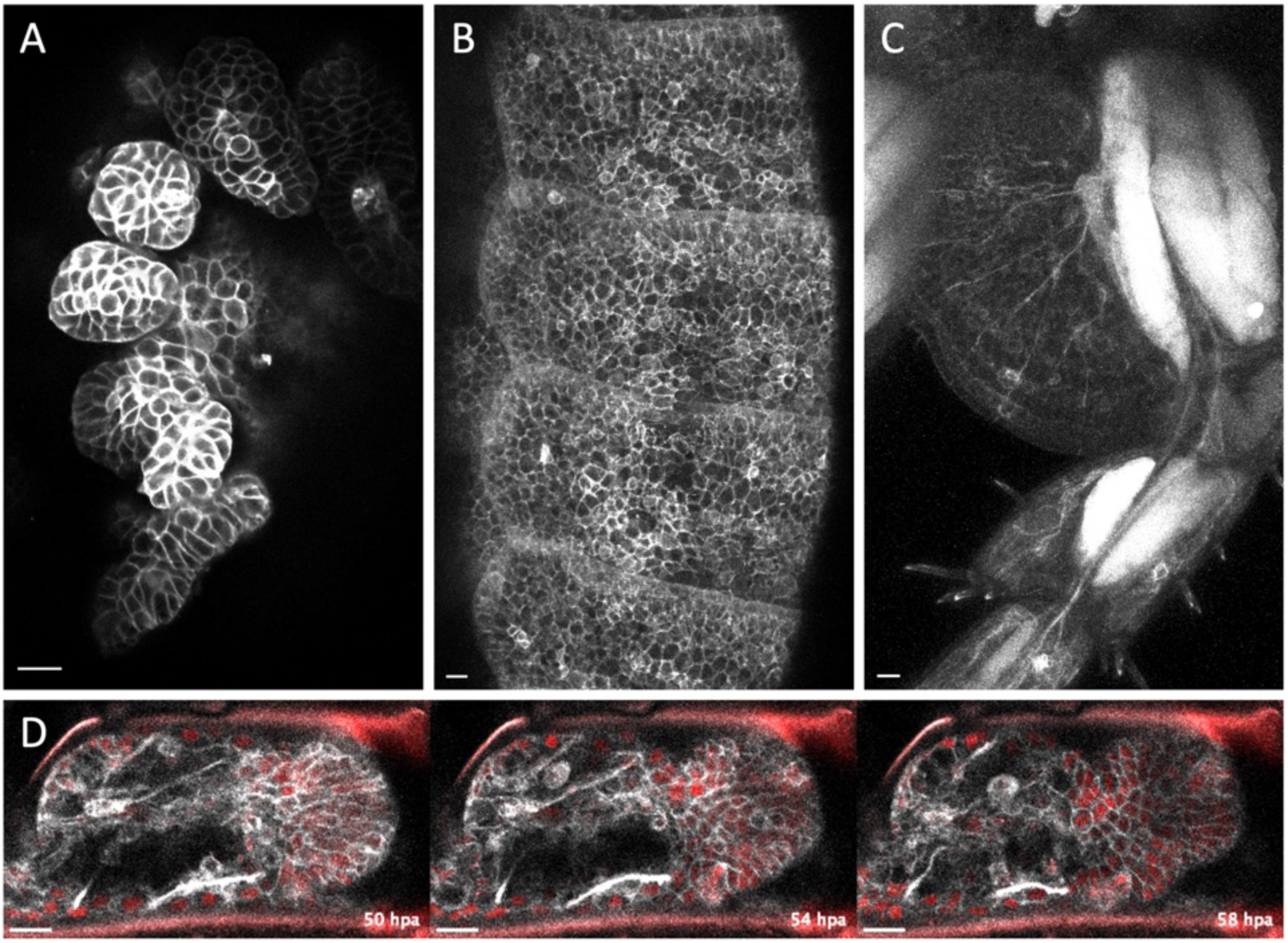
Live imaging of epidermis, muscle and neurons using selected membrane-localising tags. Live imaging of transgenic *Parhyale* expressing the mNeonGreen fluorescent protein fused with the KRas (#2) or the Src64B (#7) tags (see Methods). (A) Embryo expressing Src64B-mNeonGreen in the developing legs. (B) One-month old genetic mosaic expressing Src64B-mNeonGreen in the epidermis of trunk segments. (C) Adult genetic mosaic expressing mNeonGreen-KRas strongly in muscles and more weakly in the epidermis and neurons in the proximal part of a leg. (D) Live imaging of an adult regenerating leg expressing Src64B-mNeonGreen (in grey) and Histone2B-mRFP (labelling nuclei, in red), 50 to 58 hours post amputation (hpa). Scale bars, 20 µm.

### Outlook

We envisage that this toolkit and the results presented here will benefit researchers working in a broad range of organisms and cell types, helping to identify tags that are effective in different experimental systems. We expect it will be especially valuable in the evo-devo community, enabling the labeling and tracking of cells in a wide range of non-conventional model organisms, where tools for membrane labeling have previously been limited or unavailable. The injection of *in vitro* synthesized mRNA into early embryos offers a simple and broadly applicable method for delivering these tools in multiple species. In specific cases, it may also be possible to deliver these constructs by other means, such as transfection, electroporation or biolistics.

In organisms where transgenic approaches are already established – as in *Parhyale*, *Tribolium*, *Caenorhabditis, Macrostomum*, *Clytia* and *Nematostella* – this kit will facilitate the generation of stable transgenic reporters for studying cell morphology, cell behaviour and neural circuits during later developmental stages. Some of the tags could be used in comparative cell biology, enabling studies of membrane trafficking, lipid composition and dynamics in an evolutionary context. Finally, the toolkit’s versatility and ease-of-use could make it attractive as a teaching tool in comparative embryology courses, particularly when students have access to embryos from a variety of species.

This work has been carried out as an Open Science project. The toolkit was made available and an invitation to the research community was launched in the social media almost a year ago (Karapidaki et al. 2024). More than 15 research teams joined the effort to screen the toolkit in a wide range of organisms – some screens are ongoing. We will continue to collect and compare data from new species in the foreseeable future.

## Materials and Methods

### Plasmids

The membrane-localising tags for constructs #1 to #11 were selected as described earlier. These were fused with the N- or C-terminus of the mScarlet3 coding sequence (https://www.fpbase.org/protein/mscarlet3/; Gadella et al. 2023). The tagged (#1 to #11) and untagged/control (#0) constructs were flanked by a ∼110 bp 5’UTR fragment taken from the *Drosophila hsp70Bb* gene and a ∼703 bp 3’UTR fragment taken from the *Drosophila inflated* gene (Kapetanaki et al. 2002), and placed under the T7 promoter. In each of these constructs, a unique NotI restriction site was placed after the 3’ UTR to allow linearisation of the plasmid prior to *in vitro* transcription, and unique KasI and AsiSI restriction sites were placed on either side of the tagged mScarlet3 coding sequences, to facilitate subsequent subcloning. An ANNATG Kozak sequence was placed at the start codon, to ensure efficient translation (Kozak 1984).

The membrane-localising tags used in each construct are: (#1) 29 amino acids from the C-terminus of human HRas; (#2) 22 amino acids from the C-terminus of human KRas isoform B; (#3) same as in #2, replacing the stretch of 6 lysines in the polybasic domain by 6 arginines (Hancock et al. 1991); (#4) 62 amino acids from the C-terminus of human Rit; (#5) 20 amino acids from the N-terminus of mouse GAP43; (#6) 18 amino acids from the N-terminus of mouse Lyn; (#7) 85 amino acids from the N-terminus of *Drosophila* Src64B; (#8) 174 amino acids from the N-terminus of human PLC delta 1; (#9) 158 amino acids from the C-terminus of bovine Lactadherin; (#10) 38 amino acids from the N-terminus of *Drosophila* Wingless (signal peptide), 39 amino acids corresponding to the transmembrane domain of mouse CD8, and a 6 amino acid linker (Stapornwongkul et al. 2020); (#11) 24 amino acids from the N-terminus of rabbit lactase-phlorizin hydrolase/LPH (signal peptide), a 15 amino acid linker, the mScarlet3 protein, a 14 amino acid linker, and 31 amino acids from the C-terminus of human LFA-3/CD58 (Keller et al. 2001; Shelby et al. 2023). The constructs were produced by gene synthesis in the pTwist vector (Twist Bioscience), a high copy plasmid carrying an ampicillin resistance gene.

The plasmids are available from Addgene, at https://www.addgene.org/browse/article/28251646/; annotated sequences are available as Supplementary Data files (https://doi.org/10.5281/zenodo.17401843) and as ‘Supplemental Documents’ at Addgene.

### In vitro transcription

Plasmids were digested with NotI. After heat-inactivating NotI (15 min at 65°C) and plasmid purification, the plasmid was *in vitro* transcribed by the T7 RNA polymerase using the mMESSAGE mMACHINE T7 Transcription Kit (Thermo Fisher Scientific #AM1344), or the HighYield T7 ARCA mRNA Synthesis Kit (Jena Biosciences #RNT-102-L) for *Paracentrotus*. The mRNA was precipitated in (2.8M LiCl, overnight at −20°C), washed in 70% ethanol, and resuspended in DEPC-treated water at a concentration of 600 ng/µl. The mRNAs injected in *Platynereis* were polyadenylated using the PolyA-Tailing kit (Invitrogen #AM1350).

Alternatively, for *Clytia*, the constructs were amplified by PCR using the Phusion DNA polymerase (Thermo Scientific #F530L) and primers GCAATTAACCCTCACTAAAGGTACAAGAAGAGAACTCTGGGCG (including a T3 promoter) and AGAGTACACTGCAAGAAGTCGG. PCR products purified with Wizard SV PCR purification kit (Promega) was *in vitro* transcribed by the T3 RNA polymerase using the mMESSAGE mMACHINE T3 Transcription Kit (Thermo Fisher Scientific #AM1348). The mRNA was purified using the MegaClear kit (Thermo Scientific #AM1908).

### Microinjection and screening in *Phallusia mammillata*

Dechorionated unfertilised eggs were injected with mRNA at 1 μg/μl using a stereomicroscope, as described previously (Yasuo and McDougall 2018). Following fertilisation, eggs were cultured in plastic dishes covered with 1% agarose in filtered natural seawater at 18°C. When the embryos started gastrulating, they were transferred and oriented in transparent circular microwells made from home-made PDMS pillar arrays in a glass bottom petri dish (Godard et al. 2020). Confocal imaging of the embryos was performed on an inverted Leica Stellaris 5 equipped with a HC PL APO CS2 40x/1.10 water-immersion objective. Images were obtained in a 512 x 512 format with a 1.5x zoom factor, 2.0 Airy units, and a 0.76μm z-step size. Embryos expressing these reporters developed into swimming larvae with normal morphology.

### Microinjection and screening in *Paracentrotus lividus*

Sea urchin gametes were obtained by vigorously shaking *Paracentrotus lividus* adults. Eggs were de-jellied in acidic sea water (pH 4.5), placed on plates treated with 1% protamine sulfate and then fertilized. One-cell stage embryos were microinjected with 2 pL of microinjection solution, consisting of 300 ng/µl of mRNA encoding each membrane tag construct, 120 mM KCl, and DEPC-treated water. At least two batches, each consisting of approximately 100 embryos, were injected per construct. Injected zygotes were then washed twice with filtered sea water and incubated at 15°C overnight. The day after, embryos were transferred to 4-well plates (ThermoFisher Scientific) with filtered sea water. Paraformaldehyde was added to a final concentration of 0.2% to block embryo growth at the blastula or the gastrula stage without damaging the fluorescent signal. Embryos were initially screened on a Leica DMi8 fluorescence microscope. They were observed in greater detail on a Zeiss LSM 700 confocal microscope using a LCI Plan-Neofluar 25x/0.8 Imm Korr Ph2 M27 objective in MatTek 35 mm glass-bottomed dishes (14 mm glass, MatTek #P35G-1.5-14-C). For each construct, 4 to 10 embryos displaying positive labelling were imaged. One representative dataset per construct was selected and processed using Fiji (Schindelin et al. 2012). For the embryos shown in Figure 3, #0, #2, #6, #8, #9, #10, #11 were imaged with a laser power of 20%, #1, #3 at 15%, #4 with 100%, #5 with 25% and #7 with 7%. Embryos injected with #4 did not show any signal at any developmental stage; embryos injected with #7 started expressing the reporter at the gastrula stage.

One batch of embryos (including uninjected controls) showed abnormal development, probably due to polyspermy. In this batch, some constructs that failed to give signal at the blastula stage in other batches of embryos (#3, #7, #8, #9) showed fluorescence localised to the plasma membrane (Suppl. Figure S14); all the images for the abnormal blastulae were acquired at 20% laser power.

### Microinjection and screening in *Parhyale hawaiensis*

One- and two-cell stage embryos of *Parhyale hawaiensis* (Paris et al. 2022) were microinjected with 480 ng/µl of mRNA, as described previously (Kontarakis and Pavlopoulos 2014). The embryos were allowed to recover for approximately 24 h in filtered artificial seawater (FASW, specific gravity 1.022) at 26°C. Approximately 100 embryos were injected per construct in each round of experiments. Of these, 70 to 80% survived and a large number showed fluorescence a day after injection. Surviving embryos were initially screened on a Leica MZ16F fluorescence stereoscope. For confocal microscopy, live embryos were placed in 35mm Ibidi glass bottom dishes (Ibidi µ-Dish 35 mm, #81158) in FASW and imaged on a Zeiss LSM 800 confocal microscope using a Plan-Apochromat 20x/0.8 M27 objective (Zeiss 420650-9901-000). For live imaging (Video S1), the embryos were immobilised in the same glass bottom dishes in 3-6% methylcellulose in FASW, and imaged at 25°C.

For light sheet microscopy, embryos were placed in melted 1% low-melt agarose (Sigma-Aldrich #A9045) made in FASW. The melted agarose containing with the embryos was drawn into a glass capillary (Transferpettor caps 10µl, Brand #701902). Once the agarose had solidified, the agarose plug with the embryo was extruded from the capillary, mounted and imaged in FASW on the Zeiss LightSheet 7 microscope using a W Plan-Apochromat 20x/1.0 DIC M27 objective. Each embryo was imaged from four different views (0°, 90°, 180° and 270°) using the Zen 3.1 Black edition LS acquisition software. The Zen Blue 3.7 light sheet processing software was used to perform interactive registration, to manually align the 4 different views in x, y and z. This method generates a 3D reconstruction of the embryo in a single z-stack.

### Microinjection and screening in *Tribolium castaneum*

Microinjections were carried out following an established protocol (Berghammer et al. 2009). Adult beetles of the San Bernadino strain were transferred to boxes containing fresh white flour (type 405) and maintained at 32°C and 40% humidity. Embryos were collected after 1 hour and then kept for 1 hour at room temperature for further development. The embryos were then briefly dechorionated in 1% bleach, washed and transferred onto a microscope slide (Fisher, 0.8-1.0 mm), where they were aligned using a fine brush. The embryos were microinjected laterally close to the posterior pole, with mRNA at a concentration of 500 ng/µL. Approximately 60 embryos were injected per construct. After injection, the slide carrying the embryos was placed on agar in a humid chamber at 32°C. After 14 hours of further development, the embryos were observed on a Zeiss LSM 980 confocal microscope using a Plan-Apochromat 20X/0,8 M27 and a Plan-Apochromat 40X/1,4 Oil DIC M27 objective. The images in Suppl. Figure S3 show representative embryos injected with each construct. The images were acquired using similar settings, with only minor adjustments in the detector gain (±2% for the 20X objective and ±7% for the 40X objective) to prevent overexposure.

### Microinjection and screening in *Caenorhabditis elegans*

In vitro transcribed mRNA was injected at a concentration of 400 ng/µl into the syncytial gonad of young adult *C. elegans* hermaphrodites, following established procedures (Evans 2006). Animals were allowed to recover on nematode growth medium (NGM) plates with *E. coli* OP50. 7-8 hours after injection, injected *C. elegans* were dissected and their embryos imaged live using a Nikon Ti2 microscope controlled by Micro-Manager software and equipped with a 60x 1.4 NA objective, a Crest XLight V3 spinning disk head, a 555 nm laser for mScarlet3 excitation, and a Photometrics Prime95B camera. All the images were captured using the same settings. Brightness and contrast adjustments and a Gaussian blur filter were applied to individual images using Fiji (Schindelin et al. 2012).

### Microinjection and screening in *Platynereis dumerilii*

One- and two-cell stage embryos of *Platynereis dumerilii* (Özpolat et al. 2021) were microinjected with 170-600 ng/µl of mRNA encoding each membrane tag construct, together with 170-290 ng/µl of mRNA encoding H2B-GFP, as an additional nuclear marker. All the mRNAs except for #11 SP-GPI were polyadenylated. When necessary, the mRNAs were diluted in nuclease-free water. Approximately 20 embryos were injected per round, as described previously (Lauri et al. 2014), and most mRNAs were injected twice. Injected embryos containing four lipid droplets (an indicator of normal development) were selected on an Olympus SZX16 fluorescence stereoscope at approximately 5 hours post fertilisation (hpf). Three to six live embryos were embedded in 1% low-melting-point agarose in FASW, in a fluorinated ethylene propylene (FEP) tube (0.8 mm I.D./ 1.2 mm O.D). The refractive index of FEP (n = 1.34) closely matches that of water (n = 1.33), enabling imaging through the FEP tube. Time-lapse movies of embryo development (one to four embryos per construct) were recorded on a Flamingo light sheet fluorescence microscope (Power and Huisken 2019) with the sample chamber filled with FASW. The microscope was equipped with two 10x illumination lenses (CFI P-Fluor 10× W/ 0.30/ 3,50 objective lens; Nikon), one or two 16x detection lenses (CFI-75 LWD 16× W/ 0.80/ 3,00 objective lens; Nikon), acquiring one or two views separated by 180°, respectively. Images were captured at 10 or 15 minute intervals using 561 nm laser excitation with the same exposure time but different laser intensities; for the embryos shown in Suppl Figure S6, #1 was imaged at 10% laser power, #0, #5, #7 and #9 at 20%, #2, #4 and #6 at 30%, #8 at 40%, #3 at 60% and #10 and #11 at 100%. A time point showing clear protein localisation was selected and processed. The image stacks captured with the two illumination lenses were fused (per view) using Leonardo Fuse (Peng et al. 2025) or by average blending in BigStitcher (Hörl et al. 2019) in Fiji (Schindelin et al. 2012). Maximum intensity projections of selected focal planes were generated to optimally display the subcellular distribution of the protein.

To examine fluorescence in larvae, we followed the same microinjection approach; 10 to 80 zygotes were injected per round. After injection, animals were incubated in petri dishes with FASW for 2.5 to 7 days at 18°C. The larvae were fixed in 4% paraformaldehyde in PBS with 0.1% Tween 20 for 20-60 minutes, embedded in 2% low-melting-point agarose in PBS in a FEP tube (0.8 mm I.D./1.2 mm O.D.), and imaged on a Flamingo light sheet fluorescence microscope as described earlier, except that the sample chamber was filled with miliQ-H_2_O. For each construct, 4-10 larvae displaying positive labelling were imaged. Laser power was adjusted depending on the strength of the fluorescence: in Suppl. Figure S7, larvae for #1 and #6 were imaged at 10% laser power, #5, #7, #8, #9 and #10 at 20%, #3 and wt at 30%, #0 and #2 at 40% and #4 and #11 at 50%. For each construct, one representative dataset was processed to generate a fused image stack, using Fiji (Schindelin et al. 2012). The sharpest half of each image stack (corresponding to the region closest to the illumination lens) was extracted via the mask function and image calculator. Gaussian blur was applied to smooth the fusion boundary.

### Microinjection and screening in *Macrostomum lignano*

One-cell stage embryos of *Macrostomum lignano* (Wudarski et al. 2020) were microinjected with 400 ng/µl of mRNA encoding each membrane tag construct, together with 60 ng/µl of mRNA encoding H2B-mNeonGreen, as a nuclear marker. Approximately 30 embryos were injected per construct, using an AxioVert A1 inverted microscope (Carl Zeiss, Germany) equipped with a PatchMan NP2 holder and TransferMan NK2 micromanipulator (Eppendorf, Germany). Pressure was applied using a FemtoJet Express system (Eppendorf, Germany), with parameters adjusted based on the degree of mucus and debris. A PiezoXpert (Eppendorf) was used to penetrate the eggshell and plasma membrane. After injection the embryos were incubated at 20°C for 16 hours in artificial seawater, prior to screening on a fluorescence stereoscope. High-resolution imaging was performed using a Leica SP8X DLS confocal microscope to assess the subcellular localisation of the fluorescent signal.

### Microinjection and screening in *Clytia hemispherica*

The toolkit was tested at two different stages in *Clytia hemisphaerica*, embryos and early primary polyps (Houliston et al. 2022). Naturally spawned unfertilised eggs of *Clytia* were microinjected with 1000 ng/µl of mRNA (for screening at the blastula stage), or 250 ng/µl mRNA (for screening at the primary polyp stage), as described previously (Momose and Houliston 2007). At least two batches of 50-100 eggs were injected per construct. The injected mRNA solution was about 2-3% of the egg volume. Fertilisation was achieved by mixing injected eggs with sperm within 1 hour of spawning. Embryos were cultured in plastic dishes with artificial seawater (Red Sea Salt, 37 ppt) at 18-21°C, with 1:2000 diluted penicillin-streptomycin solution (Sigma #P4333) added for the primary polyp experiments. For screening in embryos at the blastula stage, embryos were mounted between coverslips at 5-6 hours post fertilisation and imaged on a Leica Stellaris 5 confocal microscope. For screening in early primary polyps, 3 days after injection, metamorphosis was induced by transferring planula larvae on a fluorinated ethylene propylene film (FEP, 150 µm thickness) with a drop of seawater containing GLWamide-6 (Lechable et al. 2020). Within a few hours of settlement, the primary polyps on the film were mounted on a glass slide and observed on a Leica Stellaris 5 confocal microscope, focusing on the epidermal layer.

### Microinjection and screening in *Pelagia noctiluca*

Mucus was removed from fertilized eggs with a brief incubation in a 2% L-Cysteine solution (pH 7.5) in natural sea water (38 ppt). Zygotes were transferred in a new plastic petri dish in Ca+/Mg+-free artificial sea water (530.5 mM NaCl, 10.7 mM KCl, 3.45 mM NaHCO_3_, 11.26 mM Na_2_SO_4_ in MilliQ H_2_0, pH=8.0). mRNA solutions (500 µg/µl) were injected in embryos at the one- or two-cell stage together with 3000 MW Dextran Alexa Fluor 488 (Invitrogen #D34682) (0.5 µg/µl). Approximately thirty eggs were injected per construct per batch; similar results were obtained from at least two independent injections. Embryos were washed in filtered natural sea water and incubated at 18°C. Gastrula-stage embryos (22-26 h post-fertilization) and planula larvae (46-50 h post-fertilization) were imaged under a coverslip in filtered sea water on a LSM Leica SP8X spectral confocal microscope, using a HC PL APO CS2 40x/1.30 oil-immersion objective and a 569 nm laser for mScarlet excitation. Acquisition settings were the same for gastrula and planula stages, except that the laser power was increased from 10% to 14% for the planula stage. Prior to mounting, planula larvae were immobilised by bisecting them along the short axis with dissection tools, which effectively stopped their swimming behavior.

### Microinjection and screening in *Nematostella vectensis*

In vitro transcribed mRNA was injected into fertilized eggs of the sea anemone *Nematostella* at a concentration of 50 ng/µl. The embryos were allowed to develop until late gastrula stage (30 hours post-fertilization) and then fixed and screened by immunofluorescence with an antibody that recognizes mScarlet3. We used immunofluorescence, rather than direct detection of mScarlet3, because at this stage, *Nematostella* embryos are highly refractive, which prevents imaging deeper than one third of the cell diameter below the surface. We could overcome this problem by fixing and clearing the embryos. The embryos were fixed for 1 hour in 4% paraformaldehyde in 1x PBS. After several washes in PTw (1x PBS with 0.1% Tween 20), the embryos were incubated in blocking solution (1x PBS with 1% BSA and 0.2% Triton X-100, with 5% added heat-inactivated sheep serum) for 2 hours at room temperature, followed by an overnight incubation at 4°C in blocking solution containing rabbit polyclonal pan-RFP antibody (diluted 1:1000; Chromotek pabr1-150). After eight 10-minute washes in PTw, the embryos were incubated again in blocking solution (as above). The embryos were then incubated with the goat anti-rabbit IgG AlexaFluor568 secondary antibody (1:1000 in blocking solution; Invitrogen A-11011) with the 4U/ml AlexaFluor488-phalloidin staining (Invitrogen A12379) overnight at 4°C. After eight 10 minute washes in PTw, the embryos were gradually infiltrated with Vectashield (Vector Labs) and imaged with a 63x glycerol immersion objective on a Leica Stellaris 5 confocal microscope focusing on the ectoderm.

### Quantification of fluorescence in the plasma membrane and cytoplasm

*Parhyale* embryos injected with each construct were imaged by confocal microscopy as described above, using the same optics and image acquisition settings for all constructs. Fluorescence intensity was quantified using ImageJ as follows: We selected specific optical sections in which membrane-localised fluorescence could be seen clearly on the basolateral membranes of cells. Background signal was measured by drawing a 20-pixel-wide line on the area surrounding the embryo and capturing the average intensity along that line using the Plot Profile function (⌘K). The fluorescence signal in cells was measured by drawing a 20-pixel-wide line across a selected cell, spanning the cell membranes on each side and the interior of the cell; the average fluorescence intensity along the length of that line was measured using the Plot Profile function (⌘K). The average background intensity was subtracted from all the values. The average of the two values at the plasma membrane was taken as a measure of fluorescence intensity at the plasma membrane. The average value at the interior of the cell (excluding the membrane signal) was taken as a measure of fluorescence intensity in the cytoplasm. To account for cell-to-cell and embryo-to-embryo variation we typically quantified 5 cells per embryo from 3 different embryos per construct. This quantification was performed only for constructs in which mScarlet3 was clearly localised on the plasma membrane. The same approach was used to quantify membrane and cytoplasmic signal intensity in *Phallusia*, *Paracentrotus*, *Macrostomum* and *Clytia* blastula stage embryos.

Note that these measurements underestimate the enrichment of mScarlet3 at the plasma membrane, because signal originating from the 5-10 nm wide plasma membrane becomes scattered over several pixels (several μm wide).

Figure 6 depicts the degree of plasma membrane localisation of each reporter, and also gives a rough estimation of observed fluorescence intensity at the membrane on a relative scale, for each species. In some species (*Phallusia*, *Paracentrotus*, *Parhyale*, *Macrostomum*), this represents the measured membrane/cytoplasm ratio and fluorescence intensity at the membrane, respectively, normalised on a scale of 0 to 10 within each species. In the other species, membrane localisation and intensity of fluorescence were assessed by eye and graded on a scale of 1 to 10.

### Membrane-tagged marker in *Salpingoeca rosetta*

The mStayGold-KRas reporter was generated by fusing a short linker peptide with the mStayGold fluorescent protein (Ivorra-Molla et al. 2024) to the KRas tag (22 amino acids from the C-terminus of human K-Ras isoform B). The coding sequence was adapted to match the codon usage of highly expressed genes in *S. rosetta*. The mRNA is expressed under an *S. rosetta* actin promoter and carries a 3’UTR from *S. rosetta* EF1 alpha. The construct was placed in a plasmid that also carries the Pac puromycin resistance gene expressed under an EF1 alpha promoter (Brunet et al. 2021). An annotated sequence of the plasmid is available at https://doi.org/10.5281/zenodo.17401843).

Transfection of this plasmid in *S. rosetta* cells was performed as described previously (Booth et al. 2018); transfected cells were selected by puromycin resistance (Brunet et al. 2021). Approximatively 80% of the puromycin resistant cells expressed mStayGold-KRas protein. For imaging, *S. rosetta* cells were plated on a 96-well plate (Ibidi #89626) coated with 10% poly-D-lysine (Sigma-Aldrich #P6407), immobilized with 0.004% paraformaldehyde (Electron Microscopy Sciences #15710) and imaged on a Stellaris 5 confocal microscope using an HC PL APO 63X/1.20 W CORR CS2 objective (Leica # 506346).

### Transgenic markers in *Parhyale hawaiensis*

The KRas (#2) or the Src64B (#7) tags were selected for building transgenic markers in *Parhyale*. A gene encoding the mNeonGreen fluorescent protein tagged with KRas or Src64B was placed under the *PhHS* heat-inducible promoter (Pavlopoulos et al. 2009) and cloned into a *Minos* transposon vector (Pavlopoulos and Averof 2005) carrying the *PhOpsin1-EGFP* transgenesis marker (Ramos et al. 2019). The constructs were generated by gene synthesis (Twist Bioscience, USA). Annotated sequences of the plasmids are available in the Supplementary Data files (https://doi.org/10.5281/zenodo.17401843).

The plasmids were microinjected in 1-cell stage embryos of the Chicago-F inbred line (Kao et al. 2016). The microinjected animals were raised and screened for green fluorescence in embryonic, juvenile and adult stages following a heat-shock, under a Zeiss AxioZoom.V16 microscope. Mosaic G0s were crossed with Chicago-F or with H2B-mRFP-expressing transgenic animals (Wolff et al. 2018) to generate stable transgenic lines. Mosaic G0s (Figure 8A-C) and fully transgenic G1s (Figure 8D) were imaged on a Zeiss LSM 800 confocal microscope. Live imaging of regeneration (Figure 8D) was performed as described previously (Çevrim et al. 2025), using an animal expressing both Src64B-mNeonGreen and H2B-mRFP (Wolff et al. 2018). Multiple optical sections were combined using the HeliconFocus software (HeliconSoft Ltd.; Figure 8A) or by maximum projection (Figure 8B,C), filtered and adjusted for brightness and contrast using Fiji (Schindelin et al. 2012).

## Data availability

The data files are available at the Zenodo public data repository, at this link: https://doi.org/10.5281/zenodo.17401843

## Acknowledgements

We thank Nicolas Minc, Alex McDougall, Jean Paul Vincent, Cyrille Alexandre, Yvon Jallais, Savvas Christoforidis and Maura Strigini for advice on membrane localisation tags. HY thanks Alex McDougall and Rémi Dumollard for *Phallusia* gametes. MHT thanks Rebecca Soliwoda for help with mRNA synthesis and Lisa Ulbrich for help with image processing. LL thanks Frédéric Sanchez for mRNA preparation. MLR and MIA thank Ennio Silvestri for advice on confocal imaging and in vitro transcription. YP and GB thank Artim Lange for initial tests with microinjection. JM and TB thank Mathilde Dura for the construct containing the linker sequence fused to mStayGold and Chantal Combredet for technical support with transfection. This research was supported by grants from the Agence Nationale de la Recherche (ANR-21-CE13-0044-01 ‘DeepLineage’, ANR-24-CE13-5353-01 ‘ERK-IEG’, ANR-22-CE92-0027 ‘DOLLI’ and ANR-19-CE13-0003 ‘MYODEVO’, to MA, HY, EH and LL, respectively), CNRS Biologie (‘Diversity of Biological Mechanisms’ to LL), the Alexander von Humboldt Foundation (to MHT, BL, JH), the Austrian Science Fund (PAT 2395824 and P36080, to GG), the Deutsche Forschungsgemeinschaft (DFG 1443/17-1 to GB), the European Union’s Marie Skłodowska-Curie programme (grant 101073238 to EB) and European Research Council (Consolidator Grant 101044989 ‘ORIGINEURO’ to PB), the U.S. National Institutes of Health (R01-GM138443 to DJD), the China Scholarship Council (grant 202104910052 to YP), core funding from the Institut Pasteur (to TB) and the Michael Sars Centre (to JC and PB), and a PhD fellowship from Stazione Zoologica Anton Dohrn (to MLR).

## Author contributions

IK and MA designed the experimental strategy; IK performed the experiments and analysed the results in *Parhyale*; MZ performed light-sheet microscopy in *Parhyale*; MK and BLB performed live imaging in *Parhyale*; MHT, BL and JH performed the experiments and analysed the results in *Platynereis*; TM, SA and EH performed the experiments and analysed the results in *Clytia*; HY performed the experiments and analysed the results in *Phallusia*; GG performed the experiments and analysed the results in *Nematostella*; CD and LL performed the experiments and analysed the results in *Pelagia*; BS, LL and CD developed the egg injection protocol in *Pelagia*; YP and GB performed the experiments and analysed the results in *Tribolium*; CP and DJD performed the experiments and analyzed the results in *Caenorhabditis*; MLR and MIA performed the experiments and analysed the results in *Paracentrotus*; MS, KU and EB performed the experiments and analysed the results in *Macrostomum*; JM and TB performed the experiments and analysed the results in *Saplingoeca*; JJC and PB generated codon-adapted mStayGold for *Salpingoeca*; MA selected the tags, designed the constructs and coordinated the project.

## Supporting Information

for Karapidaki et al. ‘Targeting the cell membrane in established and emerging model organisms’

### Supplementary Figures

**Figure S1.**
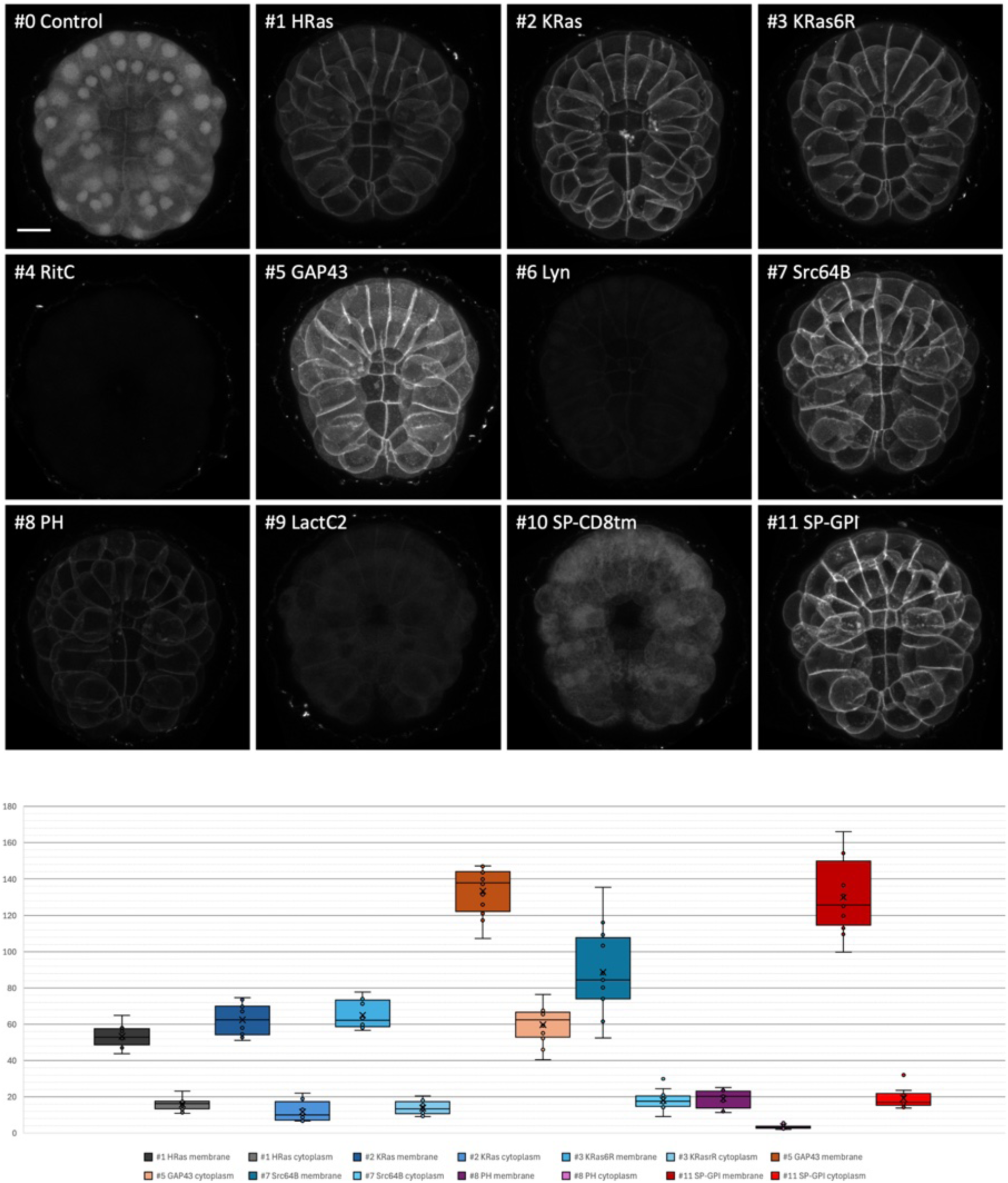
Quantification of cell membrane localisation in the tunicate *Phallusia mammillata*. (Top) Same as in Figure 2, but images were captured and displayed with the same settings, to reveal differences in fluorescence intensity. Scale bar, 25 μm. (Bottom) Quantification of fluorescence at the plasma membrane and in the cell interior, as described in the Methods.

**Figure S2.**
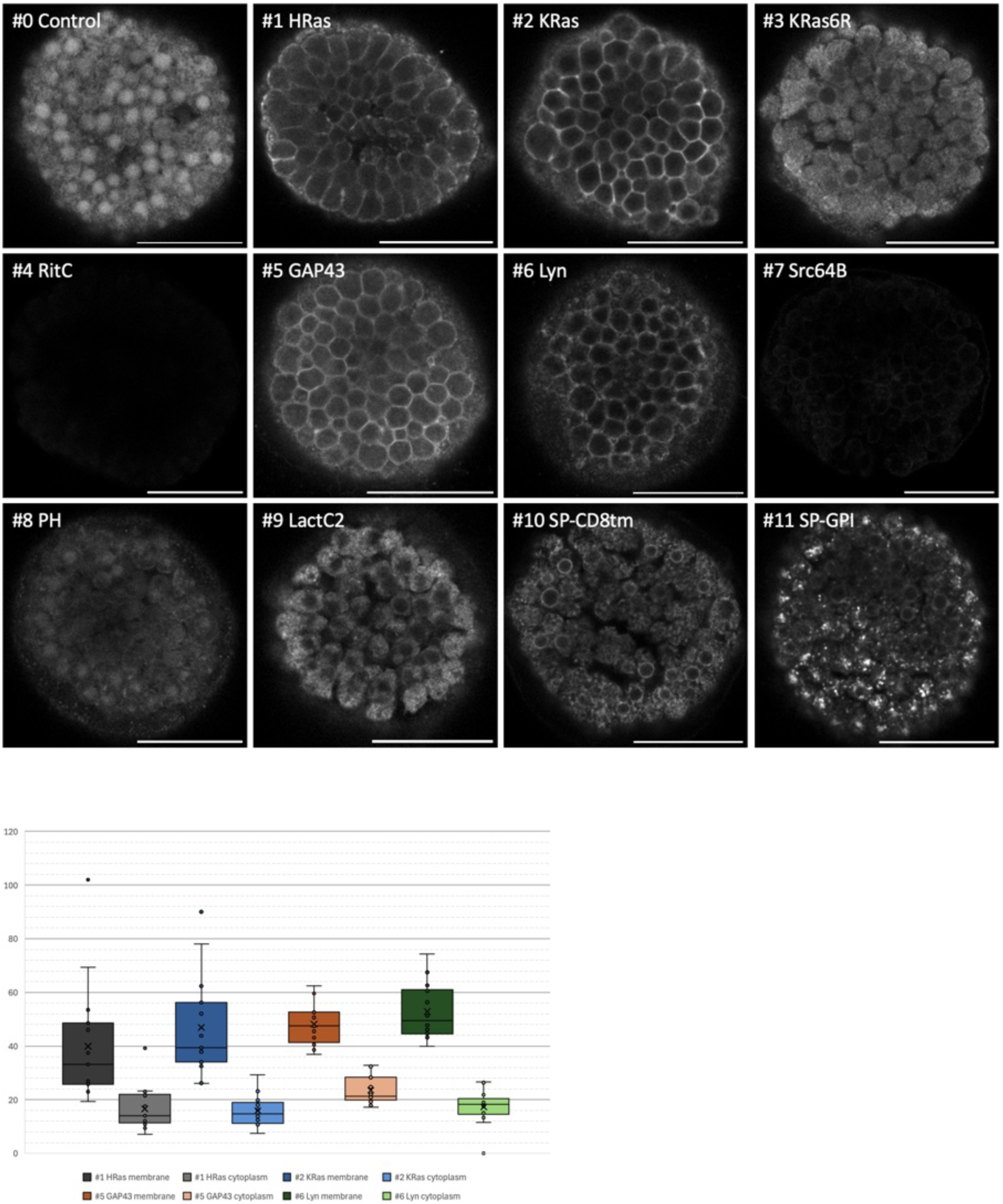
Quantification of cell membrane localisation in the sea urchin *Paracentrotus lividus*. (Top) mScarlet3 fluorescence in *Paracentrotus* blastula stage embryos. Images are single confocal planes on the embryo’s surface; unlike the images shown in Figure 3, these images were captured and displayed with the same settings to reveal differences in fluorescence intensity. Note that #7 shows no fluorescence at the blastula stage, but gives strong membrane-localised fluorescence at the gastrula stage (see Figure 3). Scale bar, 50 µm. (Bottom) Quantification of fluorescence at the plasma membrane and in the cell interior, as described in the Methods.

**Figure S3.**
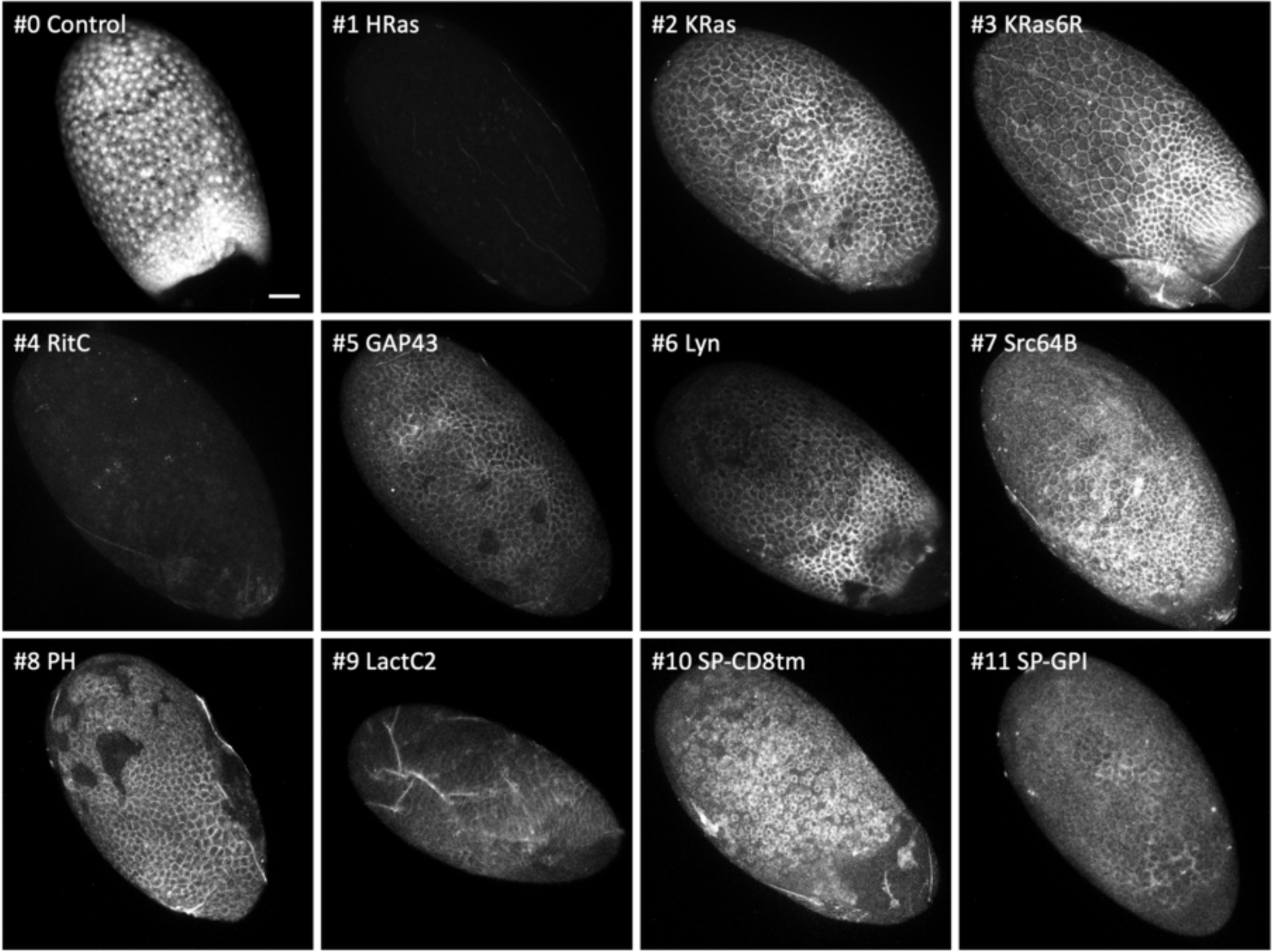
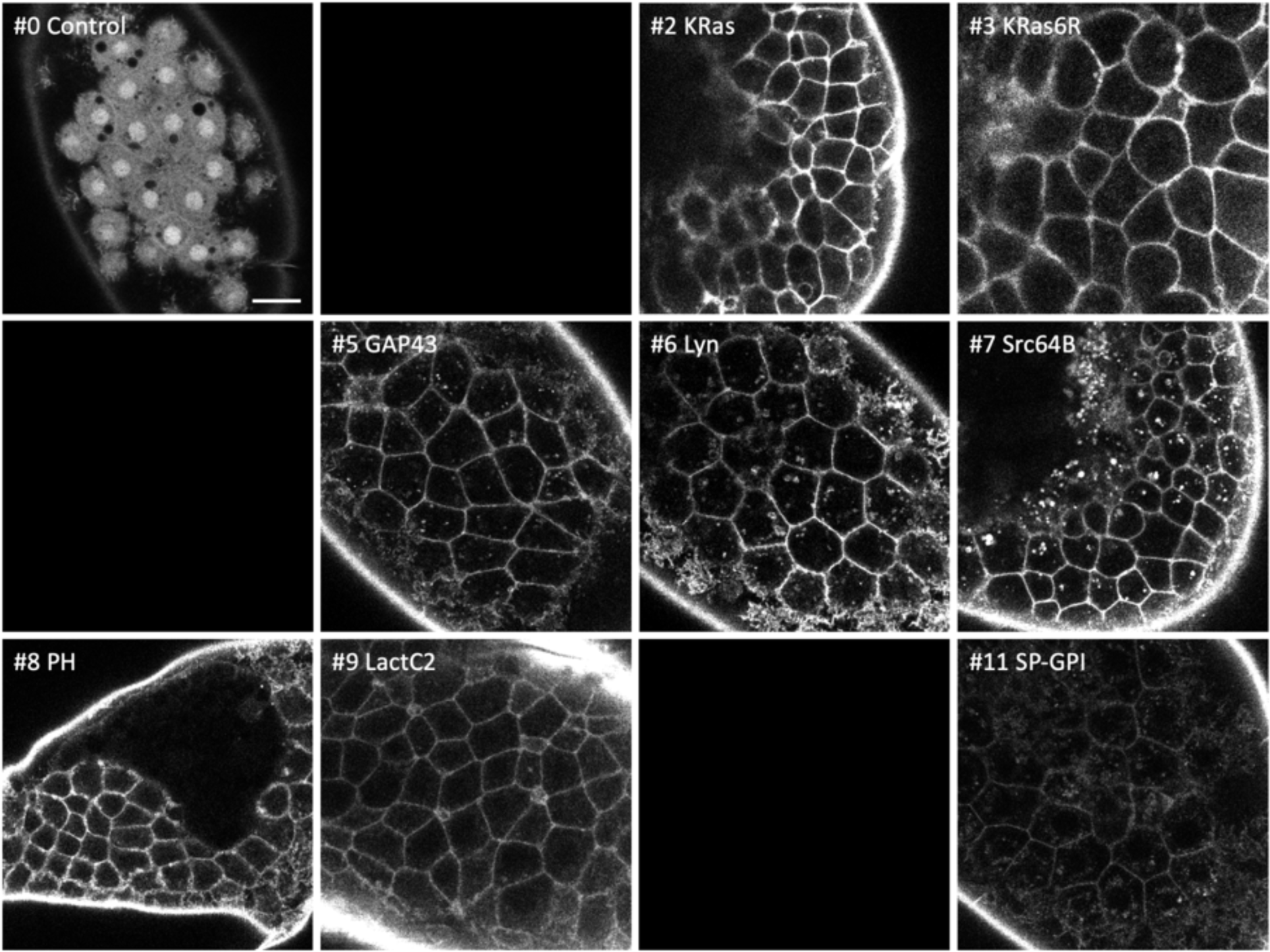
Localisation of membrane-tagged reporters in the beetle *Tribolium castaneum*. (A) mScarlet3 fluorescence in late blastoderm embryos of *Tribolium castaneum* injected with mRNAs of the membrane-tagged and control constructs. Some of the embryos show mosaic expression. Maximum intensity projections acquired with similar settings (see Methods). Scale bar, 50 μm. (B) Selected reporters imaged at higher magnification. Scale bar, 20 μm.

**Figure S4.**
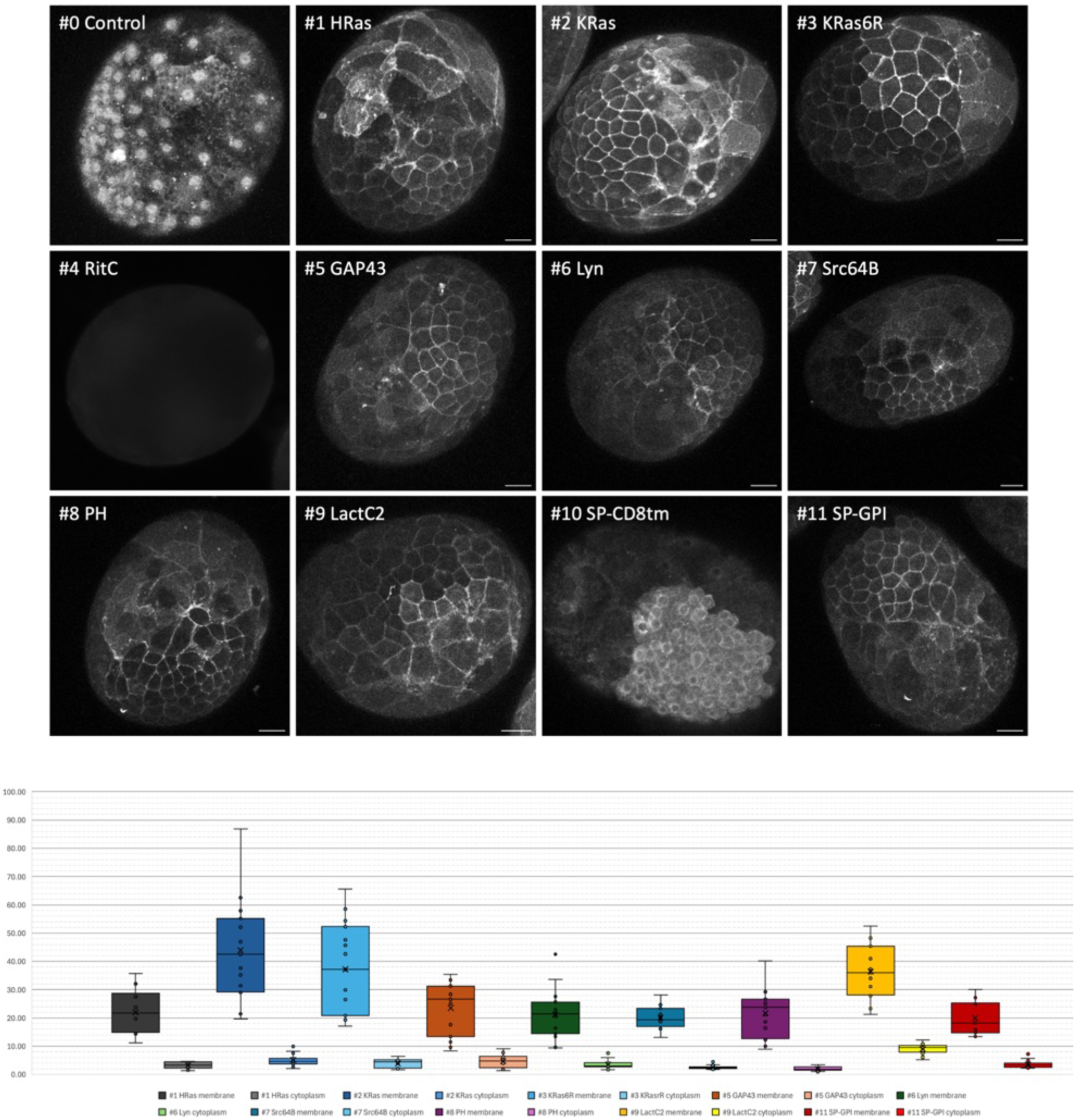
Quantification of cell membrane localisation in the crustacean *Parhyale hawaiensis*. (Top) mScarlet3 fluorescence in 1-day old *Parhyale* embryos, injected with mRNA of the membrane-tagged and control constructs. Only parts of each embryo express the reporter, due to uneven distribution of the injected mRNA. Unlike Figure 4, the images show maximum projections of images acquired by confocal microscopy, captured and displayed with the same settings to reveal differences in fluorescence intensity (except #0 and #10, which were captured in an independent experiment, and #4, which was captured by conventional fluorescence microscopy). Scale bars, 50 µm. (Bottom) Quantification of fluorescence at the plasma membrane and in the cell interior, as described in the Methods.

**Figure S5.**
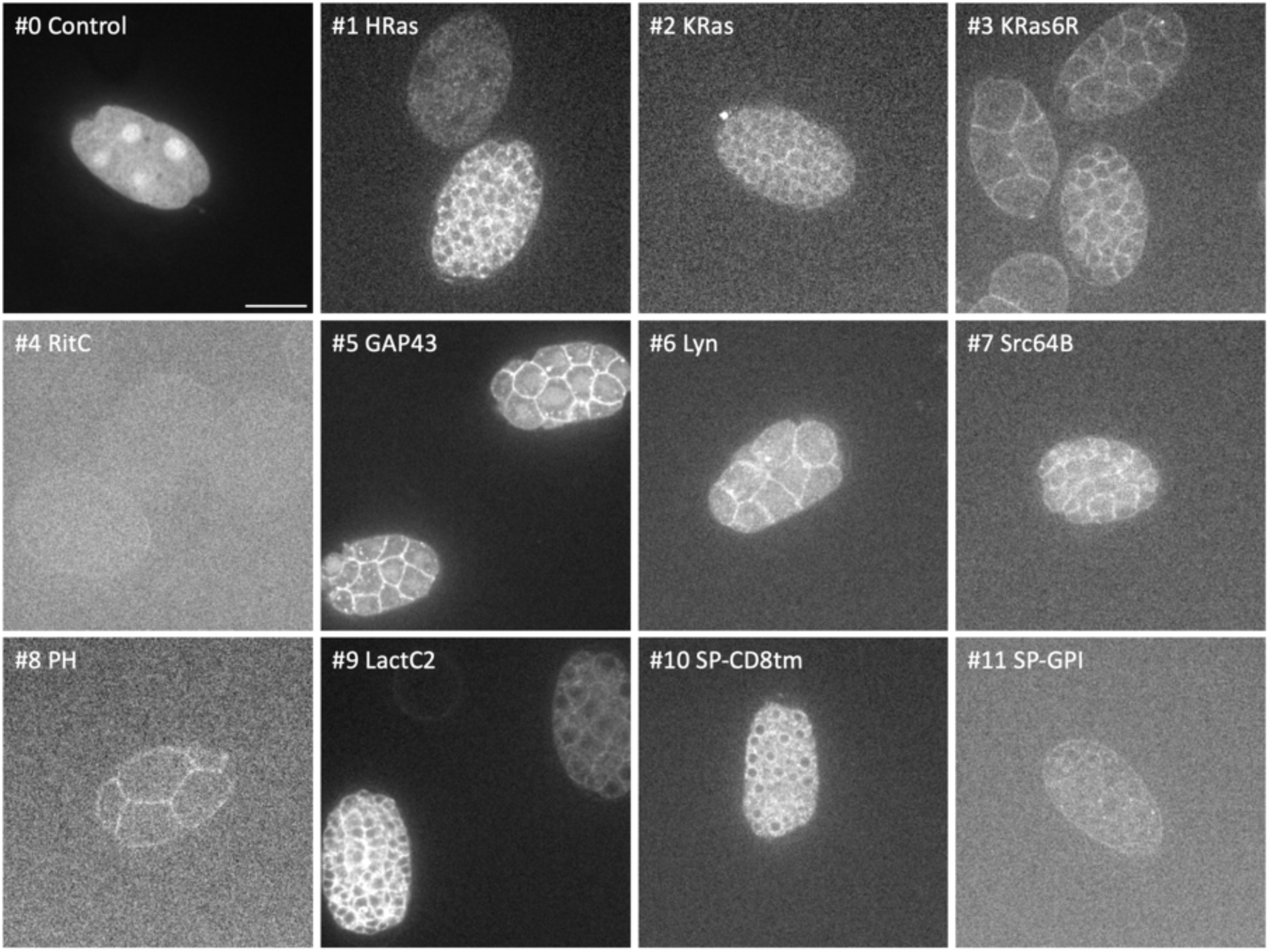
Localisation of membrane-tagged reporters in the nematode *Caenorhabditis elegans*. mScarlet3 fluorescence in early embryos of *Caenorhabditis elegans* after injecting the mRNAs of the membrane-tagged and control constructs in the syncytial gonads of their parents. The images were acquired using the same settings, but brightness and contrast were adjusted to reveal weak fluorescence. Scale bar, 20 µm.

**Figure S6.**
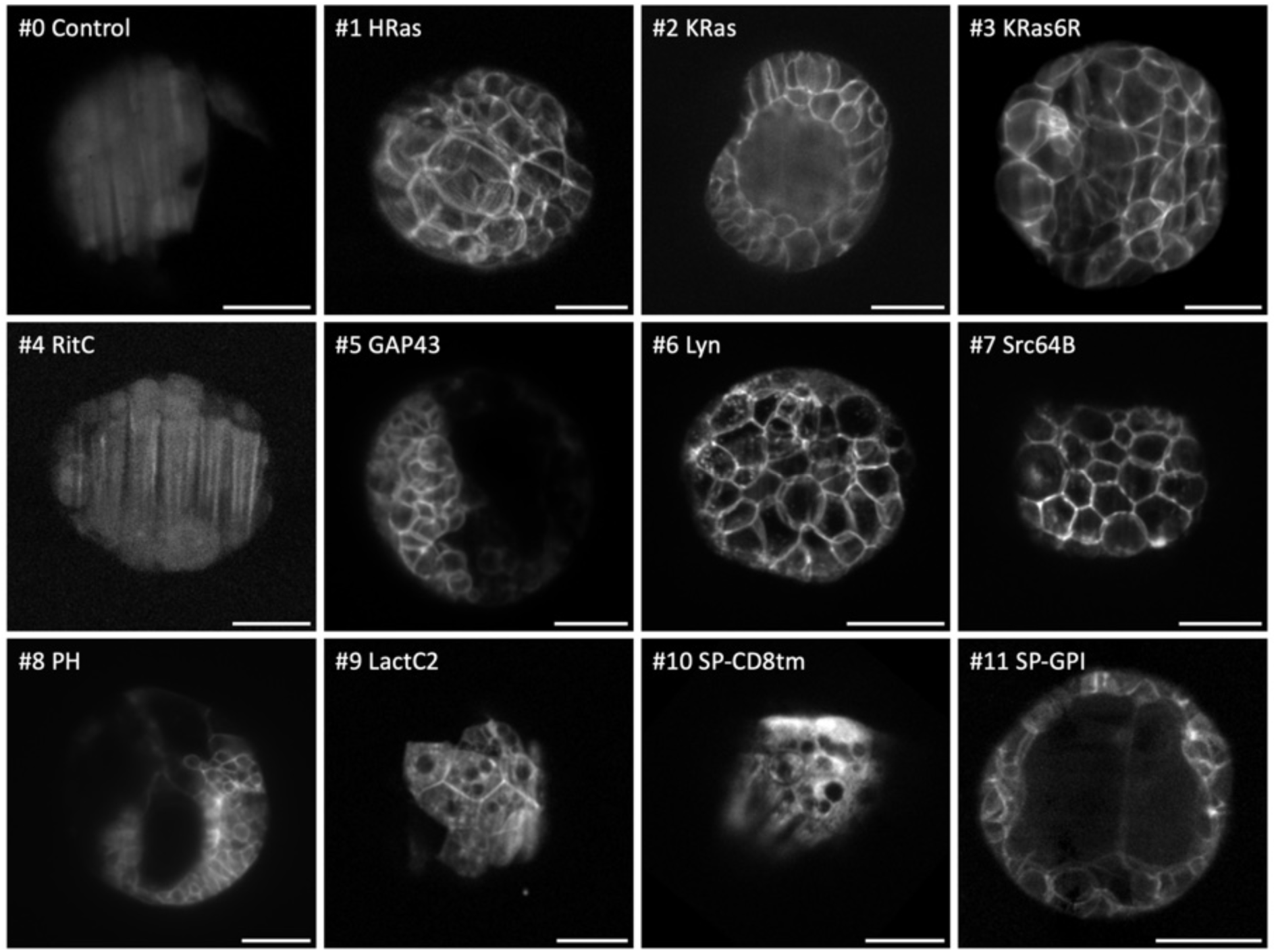
Localisation of membrane-tagged reporters in annelid *Platynereis dumerilii* embryos. mScarlet3 fluorescence in *Platynereis* embryos injected with mRNA at the 1- or 2-cell stage. Images show maximum intensity projections from light sheet microscopy, captured with different laser intensities (see Methods), at various embryonic stages: #9 at 6 hpf; #0, 1, 3, 10, 11 at 9 hpf; #7 at 10 hpf; #2, 4, 6 at 11.5 hpf; #5 at 14.5 hpf, #8 at 23 hpf (hpf, hours post fertilisation). Some embryos are mosaic because injections were performed at the 2-cell stage. Scale bar, 50 μm.

**Figure S7.**
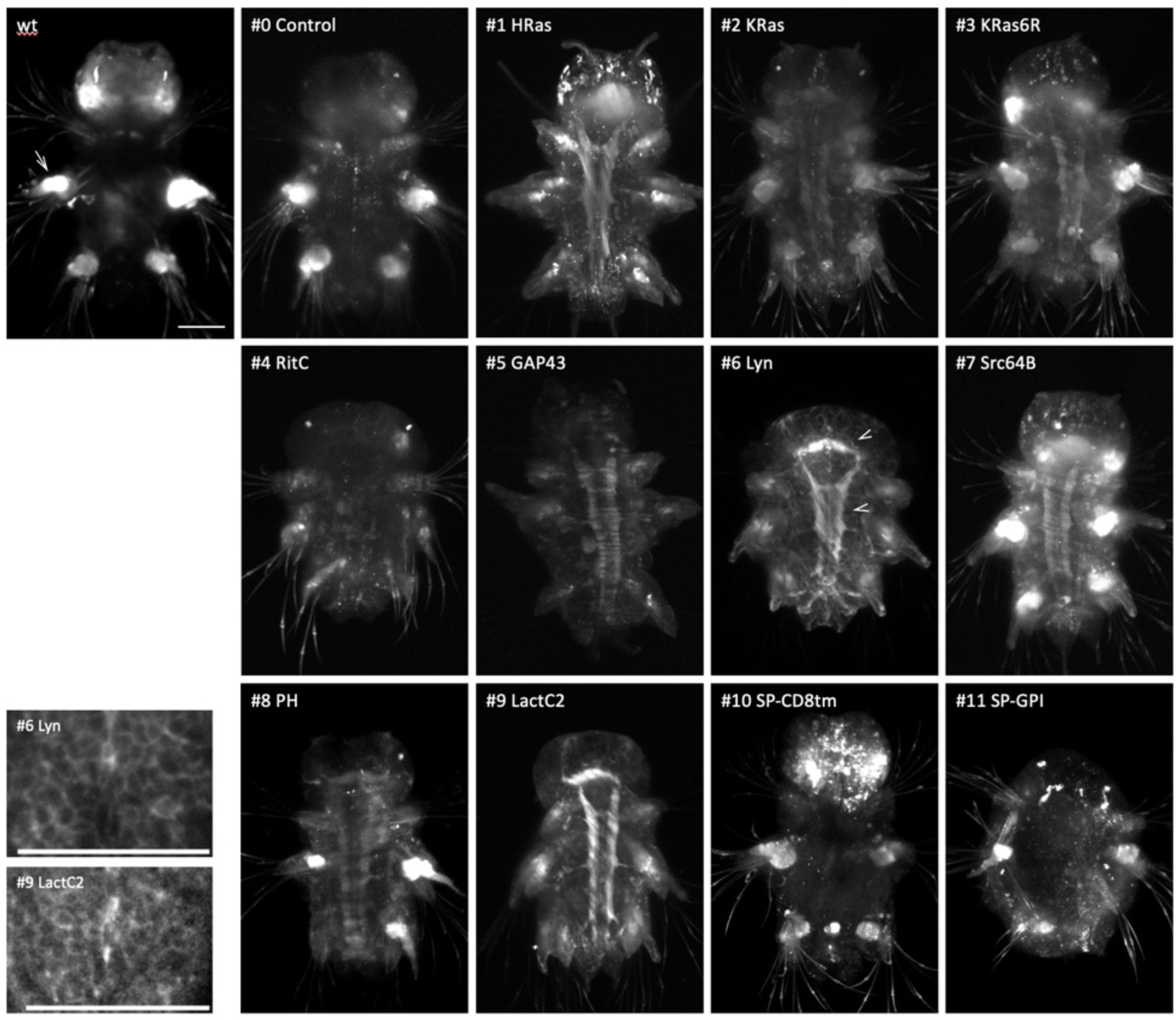
Localisation of membrane-tagged reporters in annelid *Platynereis dumerilii* larvae. mScarlet3 fluorescence in *Platynereis* larvae injected with mRNA of the membrane-tagged reporters at the 1 cell stage, compared with an uninjected larva (wt). Images show maximum intensity projections of the ventral half of each larva imaged by light sheet microscopy (see Methods), between 2.5 and 7 days post fertilisation (#6, #9 and #11 at 2.5 dpf; #3 and #4 at 3dpf; wt, #0, #2, #7, #8 and #10 at 4 dpf; #5 at 6 dpf; #1 at 7 dpf). Autofluorescence can be seen in the parapodial glands (as shown by an arrow in wt). Reporters #1 HRas, #2 KRas, #3 KRas6R, #5 GAP43, #6 Lyn, #7 Scr64B, #8 PH and #9 LactC2 showed fluorescence in the brain and ventral nerve cord (marked by arrowheads in #6). Membrane labelling outside of the nervous system was particularly visible in larvae injected with reporters #6 and #9 (higher magnification images shown in the lower left corner). Reporters #0, #4, #10 and #11 did not show a clear fluorescent signal. With some reporters (particularly #11), injecting large amounts of mRNA delayed development. The anterior of larvae is up, ventral views. Scale bars, 50 μm.

**Figure S8.**
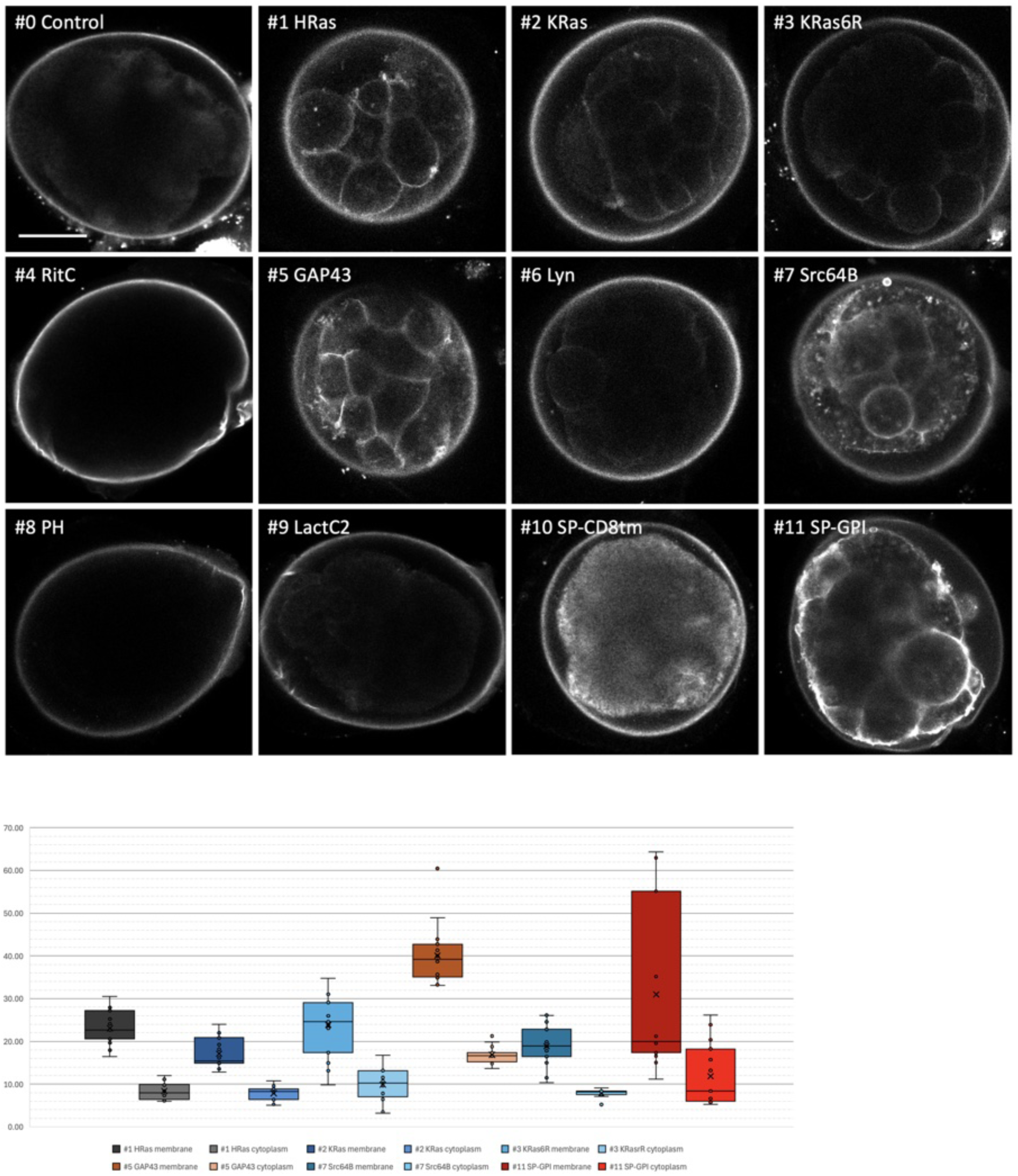
Localisation of membrane-tagged reporters in the flatworm *Macrostomum lignano*. mScarlet3 fluorescence in *Macrostomum* embryos injected with mRNA at the 1-cell stage and imaged 16 hours post injection. Images show confocal optical sections through the embryos, acquired using the same settings. Scale bar, 25 µm.

**Figure S9.**
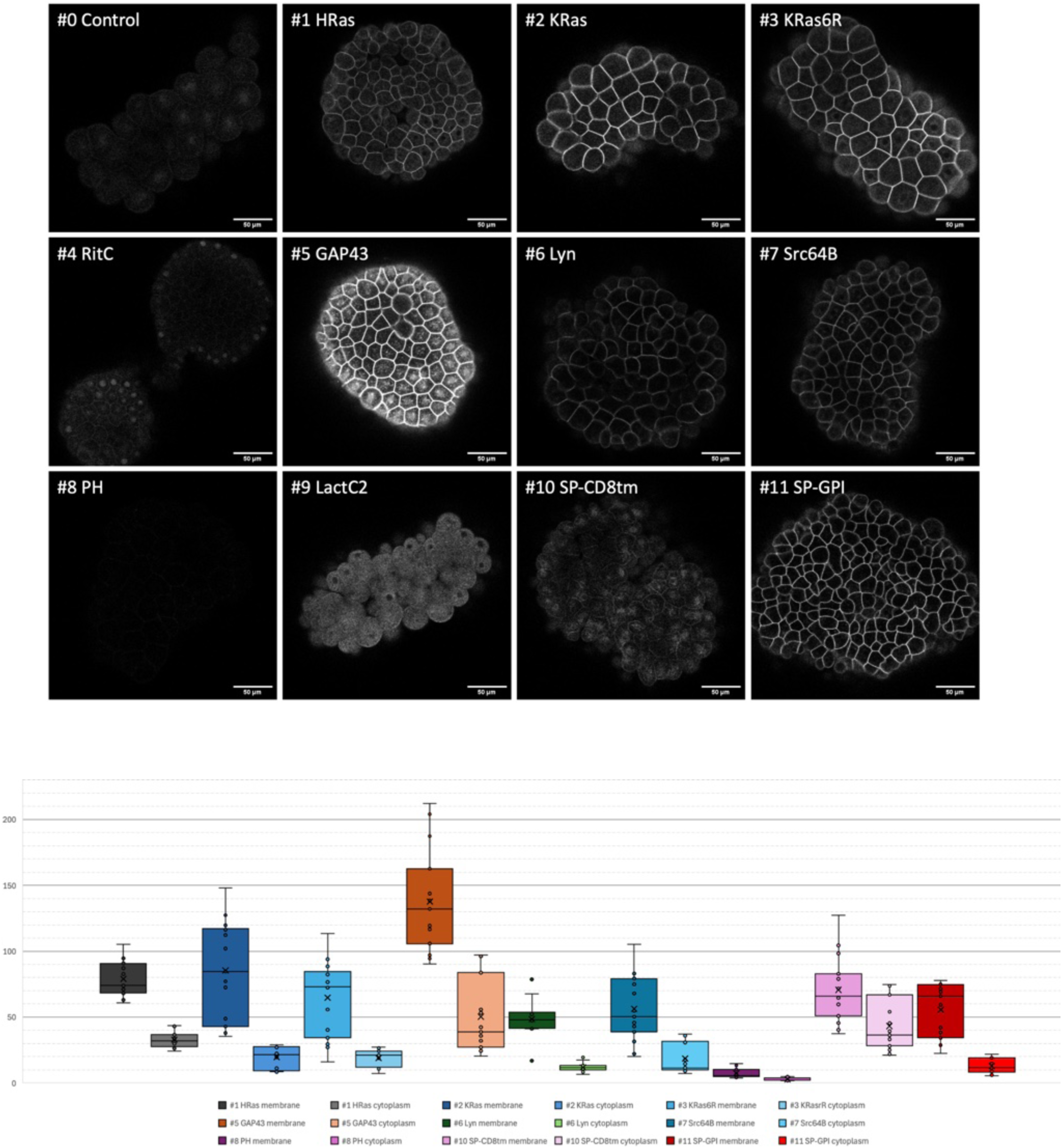
Quantification of cell membrane localisation in hydrozoan *Clytia hemisphaerica* embryos. (Top) Same as in Figure 5, but images were captured and displayed with the same settings, to reveal differences in fluorescence intensity. Scale bar, 50 µm. (Bottom) Quantification of fluorescence at the plasma membrane and in the cell interior, as described in the Methods.

**Figure S10.**
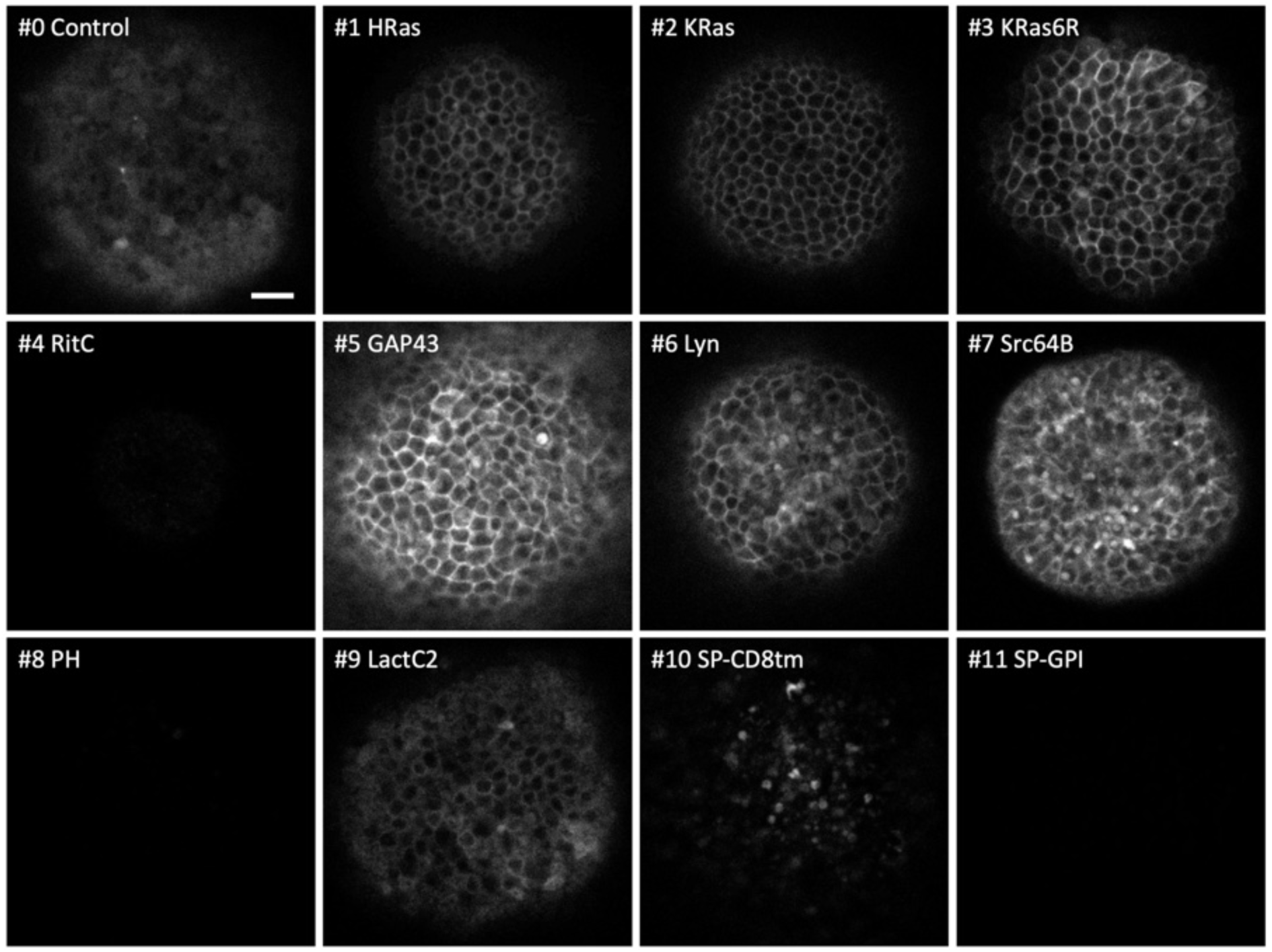
Localisation of membrane-tagged reporters in hydrozoan *Clytia hemisphaerica* polyps. mScarlet3 fluorescence in *Clytia* primary polyps a few hours after settlement, 3 days after mRNA injection into oocytes. The images are from a single confocal plane capturing the outer epidermal layer of the polyp, on the oral side. They were acquired with the same settings, so fluorescent intensities are comparable across panels. Scale bar, 10 µm.

**Figure S11.**
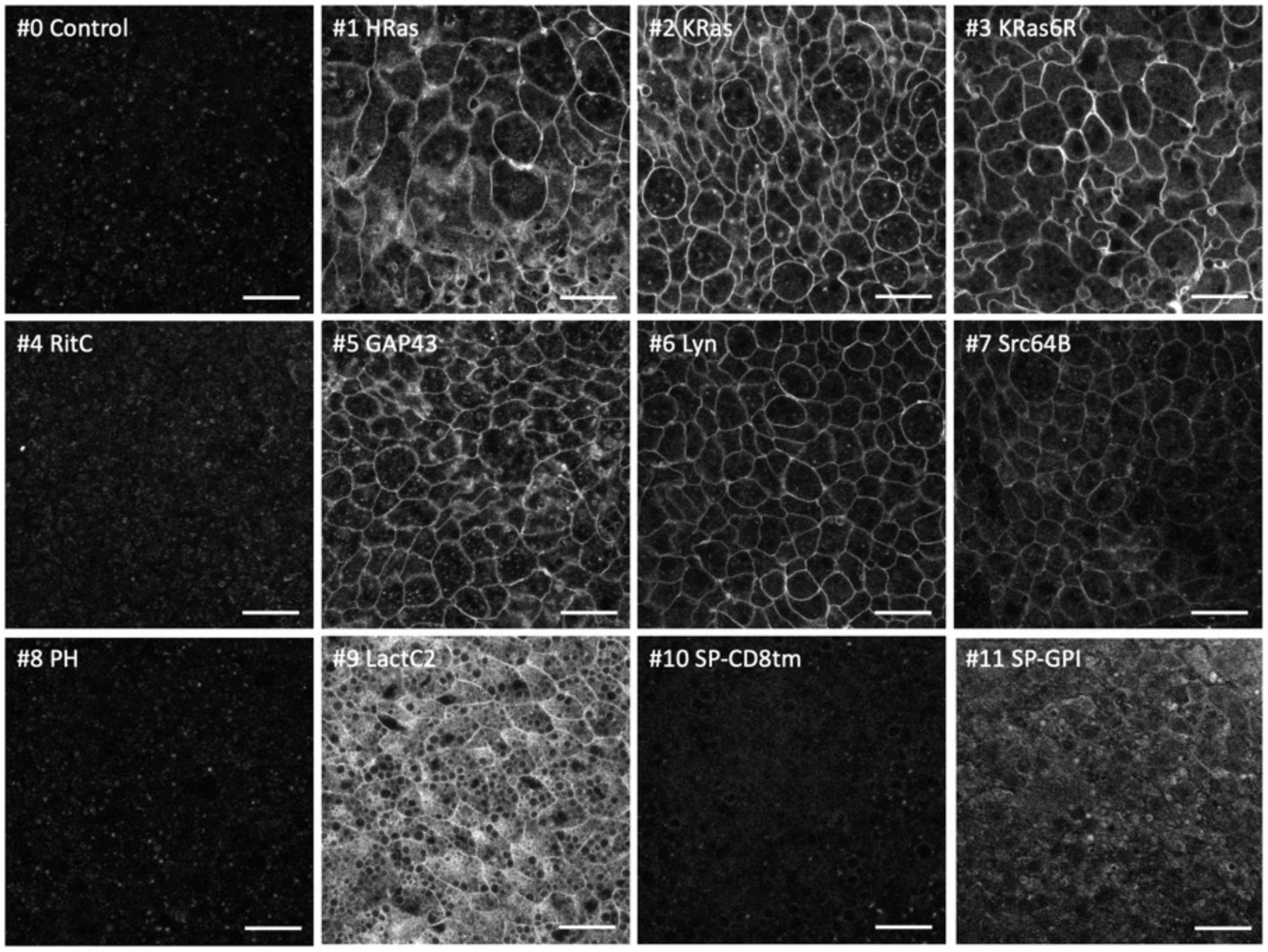
Localisation of membrane-tagged reporters in scyphozoan *Pelagia noctiluca* embryos. mScarlet fluorescence in *Pelagia noctiluca* gastrula stage embryos, 22 to 26 h after mRNA injection in zygotes. The images show a single confocal plane capturing the outer epidermal layer of the gastrula. They were captured with the same settings, so fluorescent intensities are comparable across panels. Scale bar, 20 µm.

**Figure S12.**
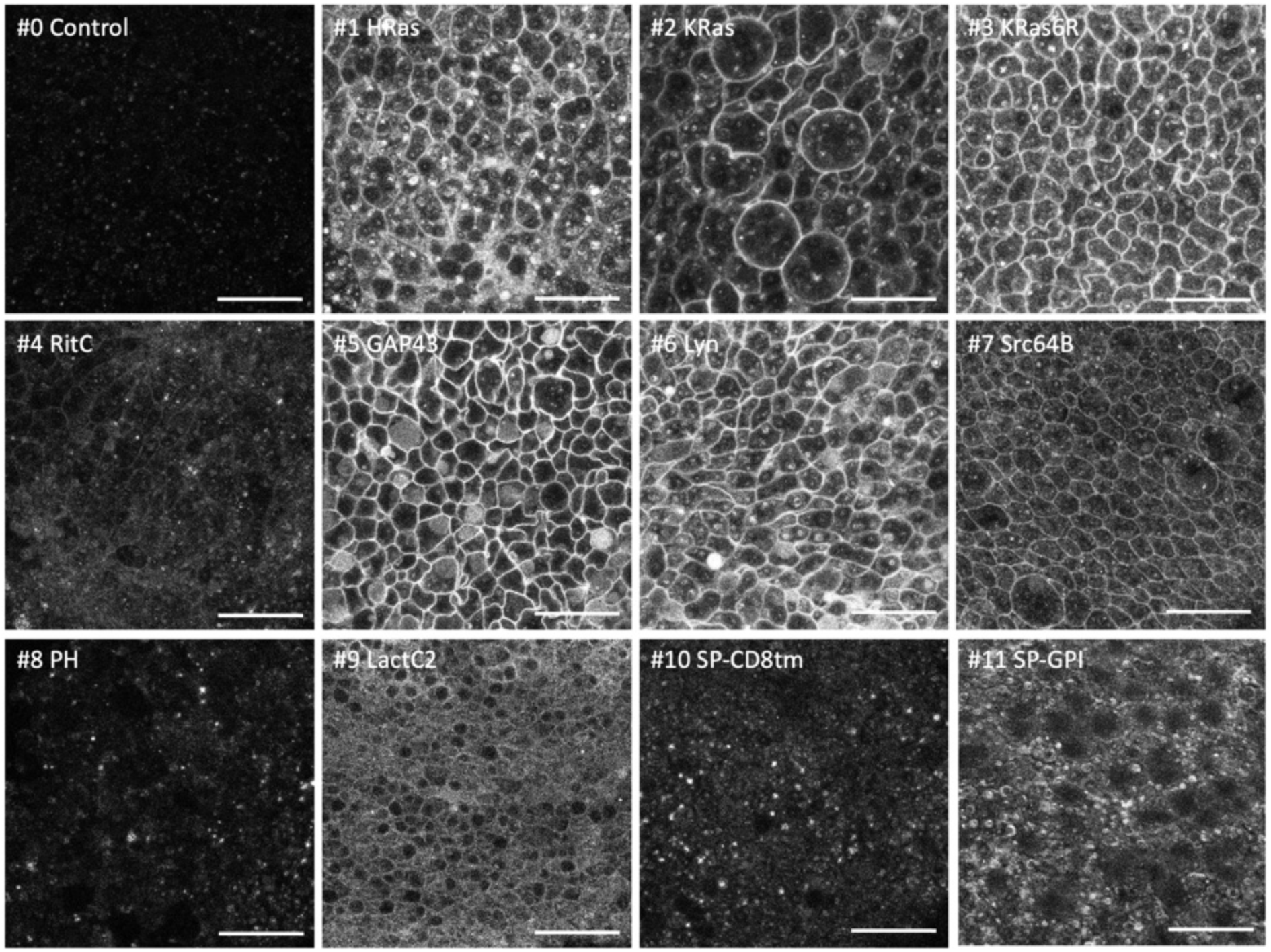
Localisation of membrane-tagged reporters in scyphozoan *Pelagia noctiluca* planulae. mScarlet3 fluorescence in *Pelagia noctiluca* planula larvae, 46 to 50 h after mRNA injection in zygotes. The images show a single confocal plane capturing the outer epidermal layer of the planula. They were captured with the same settings, so fluorescent intensities are comparable across panels. Scale bar, 20µm.

**Figure S13.**
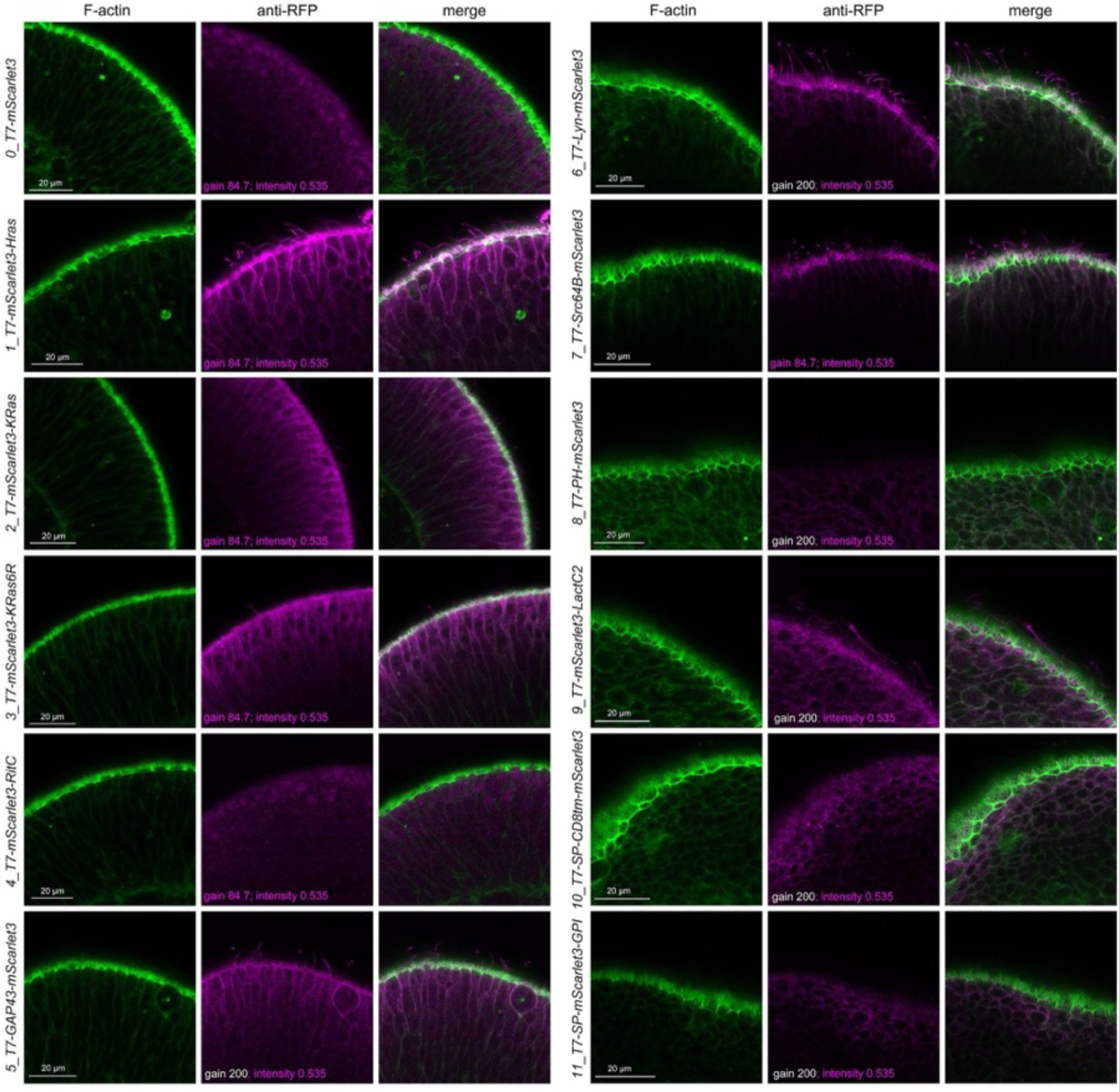
Localisation of membrane-tagged reporters in the anthozoan *Nematostella vectensis*. mScarlet3 localisation in *Nematostella* late gastrula-stage embryos, approximately 30 hours after mRNA injection. The embryos were fixed and stained with antibodies for mScarlet3 (shown in magenta) and with phalloidin (labelling actin, in green), as described in the Methods. The images show a single confocal plane across the surface of the embryo. They were acquired with the same settings, except that gain was increased for mRNAs giving weaker fluorescence (as indicated on each image). The best membrane-localising construct was #1, which gave the strongest staining of the plasma membrane, including the microvilli and cilia. Scale bars, 20 µm.

**Figure S14.**
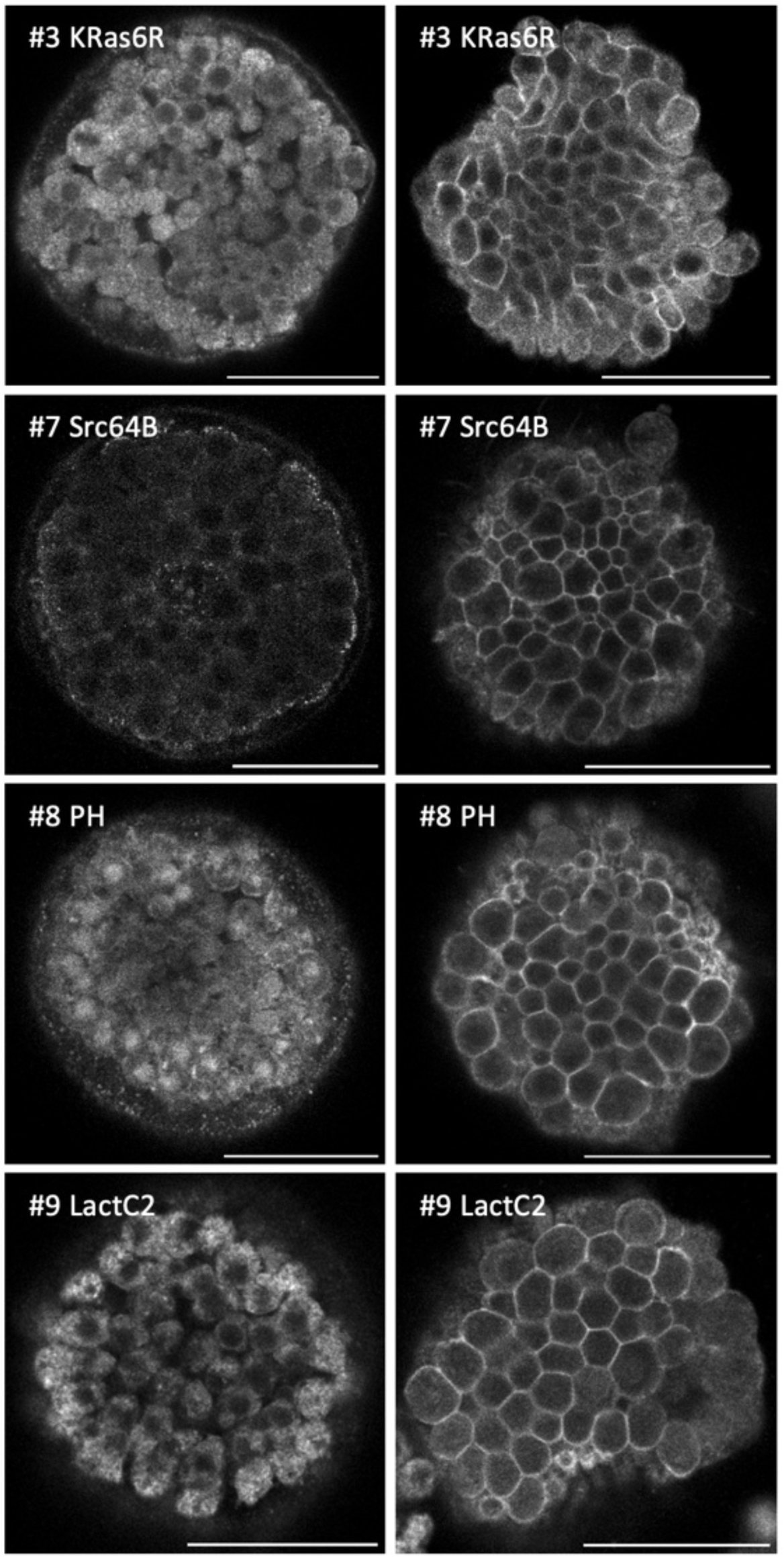
Localisation of tagged proteins in abnormal *Paracentrotus* embryos. Three reporters that did not give plasma membrane localisation of mScarlet3 in normal blastula stage embryos (#3, #8 and #9) and one that was only detected at later stages (#7), showed localisation at the plasma membrane in a batch of abnormal *Paracentrotus* blastulae, probably arising from polyspermy (see Methods). Images show single confocal planes. Normal blastulae are shown on the left, abnormal blastulae on the right. Images show single confocal planes and have been adjusted in brightness and contrast. Scale bars, 50 µm.

**Figure S15.**
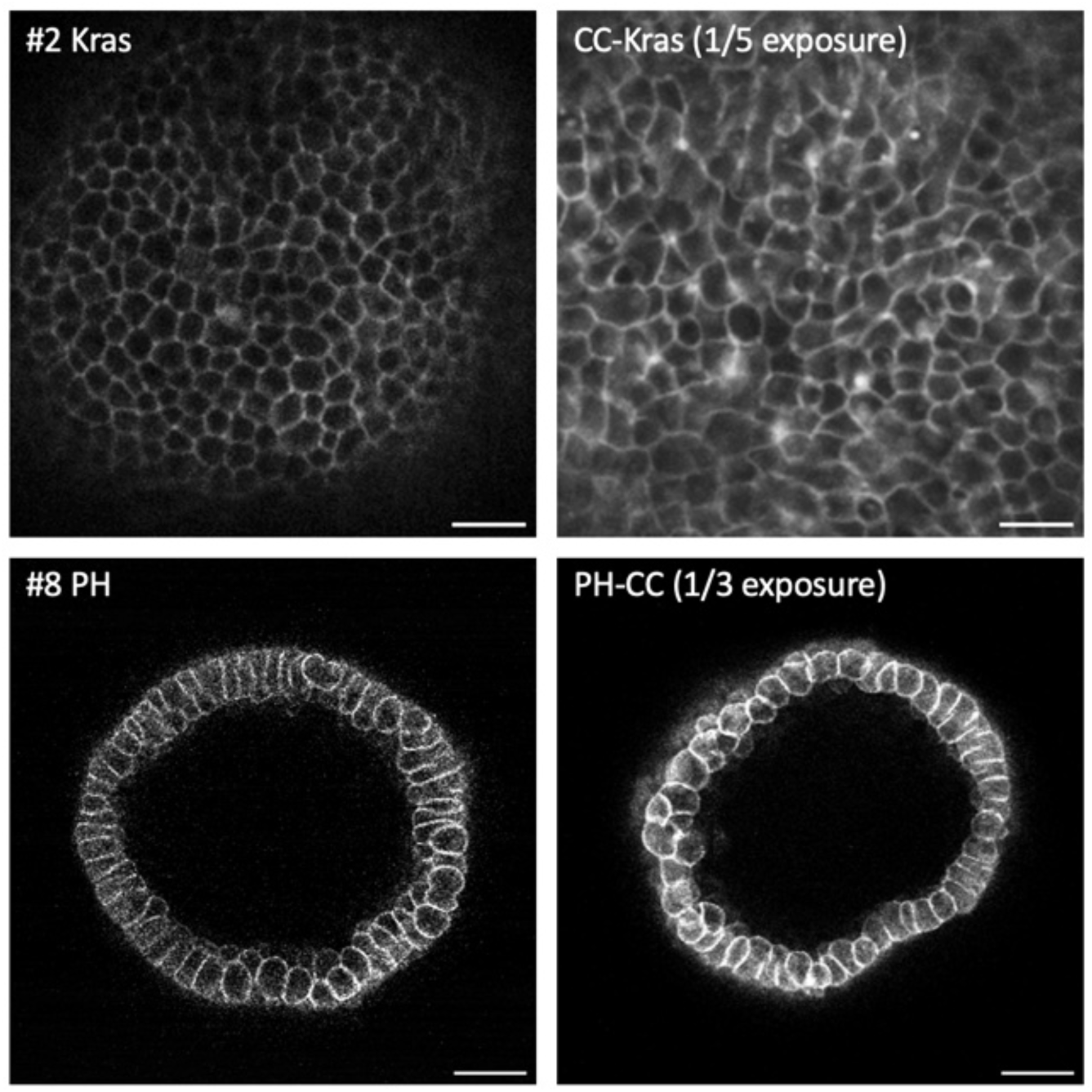
Improved reporters using endogenous UTRs and codon optimisation in *Clytia*. Using codon-optimised mCherry (adapted to the codon usage of *Clytia*, Weissbourd et al. 2021) flanked by 5’ and 3’ UTRs of a *Clytia hemispherica* genes (Uveira et al. 2024) can increase the fluorescence intensity of KRas and PH tagged reporters. (Top) *Clytia hemisphaerica* primary polyps injected with mRNA of the #2 KRas reporter (left) or codon-optimised mCherry, tagged with the same KRas tag and flanked by *Clytia* UTRs (CC-KRas, right). The CC-KRas reporter gave at least 5-fold brighter fluorescence than #2 KRas (the image was acquired with 5-fold lower exposure). Scale bars, 5 µm. (Bottom) *Clytia hemisphaerica* blastula-stage embryos injected with mRNA of the #8 PH reporter (left) or codon-optimised mCherry, fused with a codon-optimised PH tag (Uveira et al. 2024) and flanked by *Clytia* UTRs (PH-CC, right). The PH-CC reporter gave at least 3-fold brighter fluorescence than #8 PH (the image was acquired with 3-fold lower exposure). Scale bars, 50 µm.

### Supplementary Video

Video S1. Dynamics of SP-CD8tm-mScarlet3 localisation (#10) in *Parhyale* embryos

*Parhyale* embryo injected with #10 SP-CD8tm mRNA at the 1-cell stage and imaged on a confocal microscope at 20-minute time intervals. The video shows a maximum projection of multiple confocal sections, starting approximately 24 hours after injection. SP-CD8tm-mScarlet3 is partly localised on the plasma membrane at the start of the video, but it gradually shifts to a perinuclear localisation. During mitosis, SP-CD8tm-mScarlet3 can be seen aggregating in discrete regions within the cell. Different levels of fluorescence across the embryo likely reflect different amounts of mRNA inherited by each early blastomere. Scale bar, 20 µm.

The video is available at this link: https://doi.org/10.5281/zenodo.17401843

